# SnRK1 inhibits anthocyanin biosynthesis through both transcriptional regulation and direct phosphorylation and dissociation of the MYB/bHLH/TTG1 MBW complex

**DOI:** 10.1101/2022.11.21.517319

**Authors:** Ellen Broucke, Thi Tuong Vi Dang, Yi Li, Sander Hulsmans, Jelle Van Leene, Geert De Jaeger, Ildoo Hwang, Wim Van den Ende, Filip Rolland

## Abstract

Plants have evolved an extensive specialized secondary metabolism. The colorful flavonoid anthocyanins, for example, not only stimulate flower pollination and seed dispersal but also protect different tissues against high light, UV- and oxidative stress. Their biosynthesis is highly regulated by environmental and developmental cues and induced by high sucrose levels. Expression of the biosynthetic enzymes involved is controlled by a transcriptional MBW complex, comprising (R2R3) MYB- and bHLH-type transcription factors (TF) and the WD40 repeat protein TTG1. Anthocyanin biosynthesis is obviously useful but also carbon- and energy-intensive and non-vital. Consistently, the SnRK1 protein kinase, a metabolic sensor activated in carbon- and energy-depleting stress conditions, represses anthocyanin biosynthesis. Here we show that *Arabidopsis* SnRK1 represses MBW complex activity both at the transcriptional and post-translational level. In addition to repressing expression of the key transcription factor MYB75/PAP1, SnRK1 activity triggers MBW complex dissociation, associated with loss of target promoter binding, MYB75 protein degradation and nuclear export of TTG1. We also provide evidence for direct interaction with and phosphorylation of multiple MBW complex proteins. These results indicate that repression of expensive anthocyanin biosynthesis is an important strategy to save energy and redirect carbon flow to more essential processes for survival in metabolic stress conditions.

## Introduction

Plants have evolved a remarkably flexible physiology and development to cope with the constant fluctuations in their environment that affect carbon (C) and energy supplies. Photosynthesis and primary C metabolism therefore generate a variety of ‘sugar signals’ that interact with environmental and developmental cues to ensure an optimal use of resources. While sucrose has been reported to have specific regulatory effects, research has primarily focused on glucose signaling, with hexokinase as a conserved glucose sensor and the TOR kinase as a more indirect energy sensor, generally stimulating biosynthetic and growth processes (Li and Sheen, 2016). Conversely, the SNF1-related kinase1 (SnRK1) protein kinase is an evolutionarily conserved cellular fuel gauge that controls metabolism, growth, and development in response to diverse types of stress that affect photosynthesis, respiration or C allocation, while also gatekeeping important developmental transitions associated with increased energy demand or altered metabolic fluxes (Broeckx et al., 2016; Baena-González and Hanson, 2017; Baena-González and Lunn, 2020). SnRK1 generally represses biosynthetic and growth processes while activating catabolism via an extensive transcriptional reprogramming and direct phosphorylation of key metabolic enzymes (Broeckx et al., 2016). Like its fungal and animal counterparts, Sucrose Non-Fermenting1 (SNF1) and AMP-activated kinase (AMPK), SnRK1 functions as a heterotrimeric complex with a catalytic α subunit and regulatory β and γ subunits (Broeckx et al., 2016). However, while in most organisms the regulatory subunits are required for kinase activity, overexpression of the plant catalytic α subunits (encoded by SnRK1α1/KIN10 and SnRK1α2/KIN11 in *Arabidopsis*) is sufficient to confer increased SnRK1 activity in transient leaf cell assays and in transgenic plants, suggesting default activation of the plant kinase subunit (Baena-González et al., 2007; Ramon et al., 2019; Crepin and Rolland, 2019). Consistently, rather than being activated by reduced nucleotide charge, SnRK1 is inhibited by high sugar-phosphate levels, most notably by trehalose-6-P (T6P) (Zhang et al., 2009; Zhai et al., 2018), a readout and regulator of plant sucrose status (Lunn et al., 2006; Fichtner and Lunn, 2021).

Plants also evolved an extensive specialized secondary metabolism derived from primary C and nitrogen metabolism with diverse functions in establishing beneficial biotic interactions or the chemical warfare against biotic and a-biotic stressors. One important class of such metabolites are the phenolic flavonoids, including the anthocyanins that are responsible for many of the vivid red to blue colors in leaves, stems, flowers, and fruits (Grotewold, 2006). Anthocyanins are not only important in attracting animals for pollination and seed dispersal; their light absorbing qualities protect plants from high light stress and UV-damage and their accumulation has also been implicated in protecting plant tissues against pests and in oxidative and drought stress tolerance (Neill and Gould, 2003; Nakabayashi et al. 2014).

Anthocyanins are derived from the aromatic amino acid phenylalanine and from malonyl-CoA (Figure 1A). Biosynthesis involves multiple enzymes encoded by early biosynthetic genes (EBG) or late biosynthetic genes (LBG), several of which are highly regulated and intensively studied (Zhang et al., 2014). Phenylalanine ammonia-lyase (PAL), for example, converts phenylalanine into phenylpropanoids (cinnamic acid). As the first committed step of flavonoid metabolism, chalcone synthase (CHS) catalyzes the combination of the phenylpropanoid p-coumaroyl CoA with three molecules of malonyl-CoA into chalcones (tetrahydroxychalcone), after which diverse subtypes of flavonoids can be produced. While flavonol synthase (FLS) is catalyzing the production of the colorless flavonols (from dihydroflavonols), dihydroflavonol 4-reductase (DFR) and then leucoanthocyanidin dioxygenase (LDOX) activity specifically direct C fluxes toward anthocyanin biosynthesis via the subsequent glycosylation and acylation of a set of anthocyanidins (Winkel-Shirley, 1999)(Figure 1A). PAL and CHS are encoded by EBGs, while DFR, LDOX and, for example, UDP-glucose:flavonoid 3-0-glucosyl-transferase (UFGT) are encoded by LBGs. Expression of these late biosynthetic enzymes is transcriptionally regulated by a core heterotrimeric MBW complex, comprising (R2R3) MYB- and bHLH-type transcription factors (TFs) associated by the scaffolding WD40 repeat protein TRANSPARENT TESTA GLABRA1 (TTG1). Depending on the MYB and bHLH TF subunits involved, MBW complexes are also implicated in other cellular differentiation pathways, controlling epidermal cell fate (trichome and root hair cell identity) and seed coat development (Walker et al., 1999; Zhang et al., 2003; Ramsay and Glover, 2005). In the complexes regulating anthocyanin LBGs in vegetative tissues, the MYB TF subunit is encoded by *MYB75/PAP1* (*PRODUCTION OF ANTHOCYANIN PIGMENT1*) or *MYB90/PAP2* (Borevitz et al., 2000; Gonzalez et al., 2008; Deng and Lu, 2017). *GL3* (*GLABRA3*), *bHLH2/EGL3* (*ENHANCER OF GLABRA3*) or *bHLH42/TT8* encode the bHLH subunit (Zhang et al., 2003).

**Figure 1.**
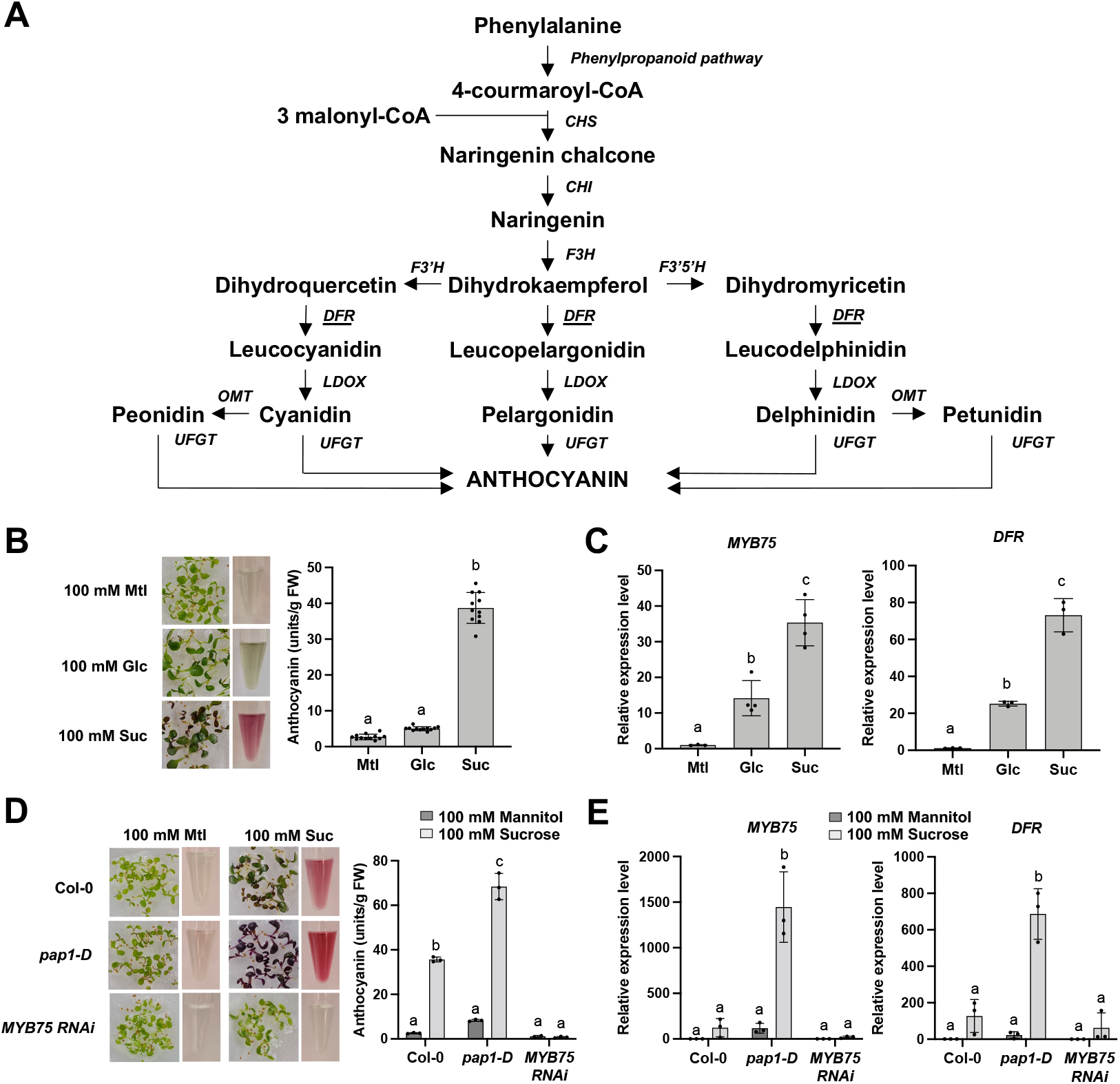
Sugar-induced anthocyanin biosynthesis is sucrose-specific. (A) Schematic representation of the anthocyanin biosynthesis pathway. Anthocyanins are synthesized from phenylalanine by both early (CHS, CHI, F3H, F3’H, and F3’5’H) and late biosynthetic enzymes (DFR, LDOX, and UFGT). CHS: chalcone synthase, CHI: chalcone isomerase, F3H: flavanone 3-hydroxylase, F3’H: flavonoid 3’-hydroxylase, F3’5’H: flavonoid 3’,5’-hydroxylase, DFR: dihydroflavonol 4-reductase, LDOX: leucoanthocyanidin dioxygenase, UFGT: UDP glucose flavonoid 3-O-glucosyltransferase. (B) Relative anthocyanin levels of 7-day-old Col-0 wildtype seedlings grown in ½ MS medium supplemented with 100 mM mannitol, 100 mM glucose or 100 mM sucrose. Values are averages with SD, n = 11 biological repeats. One-way ANOVA statistical analysis was performed in GraphPad Prism v9, letters represent statistically significant differences, p<0.0001. FW, fresh weight. (C) qRT-PCR analysis of MYB75 and DFR gene expression levels in 7-day-old Col-0 wildtype seedlings grown in ½ MS medium supplemented with 100 mM mannitol, 100 mM glucose or 100 mM sucrose. Values are averages with SD, n = 4 biological repeats. One-way ANOVA statistical analysis was performed in GraphPad Prism v9, letters represent statistically significant differences, p<0.025. (D) Relative anthocyanin levels of 7-day-old Col-0 wildtype, pap1-D and MYB75 RNAi seedlings grown in ½ MS medium supplemented with 100 mM mannitol or 100 mM sucrose. Values are averages with SD, n = 3 biological repeats. Two-way ANOVA statistical analysis was performed in GraphPad Prism v9, letters represent statistically significant differences, p<0.0001. FW, fresh weight. (E) qRT-PCR analysis of MYB75 and DFR gene expression levels in 7-day-old Col-0, pap1-D and MYB75 RNAi seedlings grown in ½ MS medium supplemented with 100 mM mannitol or 100 mM sucrose. Values are averages with SD, n = 3 biological repeats. Two-way ANOVA statistical analysis was performed in GraphPad Prism v9, letters represent statistically significant differences, p<0.0001.

The diverse functions of anthocyanins require a tight regulation of their production. Developmental and environmental cues such as light, temperature, phosphate and nitrogen status (Lea et al., 2007; Feyissa et al., 2009; Rowan et al., 2009; Hsieh et al., 2009; Gou et al., 2011; Maier et al., 2013; Liao et al., 2022) as well as hormone (Loreti et al., 2008; Jeong et al., 2010; Lewis et al., 2011) and innate immune signaling (Saijo et al., 2009; Serrano et al., 2012) are known to modulate anthocyanin biosynthesis. While the MBW complex proteins themselves (and especially the MYB TFs) are transcriptionally regulated by different cues (Shin et al., 2007; An et al., 2012; Shin et al., 2013; Meng et al. 2020; Gangappa and Botto, 2014; Li, 2015; Chang et al., 2008; Li and Zachgo, 2013; Viola et al., 2016), MBW complex activity is also controlled post-translationally. MYB75 and MYB90, for example, are targeted for degradation by COP1/SPA in the dark (Maier et al., 2013), while phosphorylation by MPK4 increases MYB75 stability (Li et al., 2016). SUMOylation by SIZ1 also stabilizes MYB75 in high light conditions (Zheng et al., 2020). GL3 and bHLH2/EGL3 are targeted for degradation by the ubiquitin ligase UPL3, affecting both anthocyanin biosynthesis and trichome development (Patra et al., 2013). MBW complex activity is also regulated by direct interaction with other TFs, including the competitively binding single MYB domain (R3-MYB) MYBL2 TF (Matsui et al., 2008; Dubos et al., 2008) and miR156-regulated SPL9 TF (Gou et al., 2011), and with the JAZ proteins, negative regulators of JA signaling (Qi et al., 2011). Interestingly, the DELLA proteins (proteasome-degraded negative regulators of gibberellin-signaling and major points of integration of different hormone signals) can sequester MYBL2 and the JAZ proteins, leading to MBW complex activation (Xie et al., 2016). In addition, a recent study related the negative regulation of anthocyanin accumulation by PIF4 to the competitive binding of PIF4 and TT8/bHLH42 to MYB75 (Qin et al., 2022).

High sucrose levels are an important trigger for attractive and/or protective anthocyanin accumulation in different species. This is most obvious in ripe fruits, sink leaves or tissues accumulating sucrose in response to, for example, high light, cold or drought stress. QTL and mutant analyses identified *MYB75* expression as a critical component in sucrose-specific induction of anthocyanin biosynthesis (Teng et al., 2005; Gonzalez et al., 2008; Li et al., 2016). Conversely, increased SnRK1 activity is known to repress *MYB75* expression (Baena-González et al., 2007) and anthocyanin biosynthesis, also consistent with the reported positive correlation of altered levels of T6P with anthocyanin accumulation during sugar-induced senescence (Wingler et al., 2012). We have previously identified sucrose-induced stabilization of the DELLA proteins as a new mechanism of *MYB75* and anthocyanin biosynthesis induction, but the DELLA proteins and T6P/SnRK1 signaling may act independently in sucrose regulation of anthocyanin biosynthesis (Li et al., 2014). More recently, anthocyanin biosynthesis during high light acclimation was also reported to require inactivation of SnRK1 by chloroplast-derived (triose-P export-dependent) sugar signals (Zirngibl et al., 2022).

How C and energy status are integrating developmental and environmental cues at the level of MBW complex activity to control anthocyanin biosynthesis is, however, still not fully understood. We therefore further explored how SnRK1, as an indirect metabolic integrator of very diverse cues, represses this important but non-vital and expensive process, thereby prioritizing and redirecting C fluxes to more essential functions for survival. We used cellular assays in combination with seedling assays and characterization of mutant and transgenic plants to provide evidence that *Arabidopsis* SnRK1 directly interferes with MBW complex formation and stability.

## Results

### SnRK1 represses sucrose-induced anthocyanin biosynthesis

We started our analyses by setting up a convenient experimental system. Growing Col-0 wildtype *Arabidopsis* seedlings in 6-well plates in ½ MS medium supplemented with different sugars confirmed the sucrose-specificity of sugar-induced anthocyanin biosynthesis (Teng et al., 2005; Solfanelli et al., 2006; Li et al., 2014). Consistent with the significantly stronger increase of anthocyanin levels after 3 days in sucrose-supplemented than after 3 days in glucose-supplemented medium (Figure 1B), sucrose also more effectively induced the expression of *MYB75/PAP1* and its *DFR* target gene 6h after supplementation (Figure 1C). Mannitol was used as an osmotic control. Genes encoding more upstream flavonoid biosynthesis enzymes (*CHS, CHI, F3H*) only showed a moderate induction upon sucrose treatment (Figure S1A), while the induction of *LDOX*, encoding the enzyme acting downstream of DFR was also very pronounced. Expression of the other *MYB* and *bHLH* TFs in the MBW complex (MYB90, bHLH42/TT8, bHLH2/EGL3 and GL3) also increased significantly upon sucrose treatment, albeit not as strongly as *MYB75*. Expression of the unique *TTG1* subunit does not appear to be regulated by sucrose supply (Figure S1B). The strong induction of *MYB75* expression is consistent with its previously suggested critical role in sucrose-induced anthocyanin biosynthesis (Teng et al., 2005). We confirmed this key role in our experimental setup by using Col-0 wildtype, *pap1-D* (*MYB75 OX*; Borevitz et al., 2000) and *MYB75 RNAi* (Jover-Gill et al., 2014) seedlings (Figure 1D). We observed a consistent significantly increased *MYB75* and *DFR* expression in *pap1-D* mutant seedlings and a significantly decreased *MYB75* and *DFR* expression in *MYB75 RNAi* seedlings. These changes in gene expression levels were seen in control conditions and especially in sucrose-rich conditions (Figure 1E). In the *pap1-D* overexpression line, containing a T-DNA with multiple 35S enhancers 3’ of the coding sequence (Borevitz et al., 2000), *MYB75* expression (transcript level) apparently is still responsive to sucrose.

Finally, we confirmed the repression of anthocyanin biosynthesis by SnRK1 activity using Ler-0 wildtype and *SnRK1α1 OX* seedlings (Baena-González *et al*., 2007). *SnRK1α1 OX* significantly reduced sucrose-induced anthocyanin biosynthesis (Figure 2A) and *MYB75* and *DFR* expression (Figure 2B). A *dfr* mutant line was included as a control, demonstrating that DFR is a key enzyme in anthocyanin biosynthesis. *SnRK1α1 RNAi* seedlings did not show significantly increased sucrose induction of anthocyanin (Figure 2A) or *MYB75* and *DFR* transcript levels (Figure 2B), probably due to the partial effect of RNAi and functional redundancy with SnRK1α2 (Wilson et al., 2011). Another conceivable explanation is that no additional effect upon lowering SnRK1 transcripts could be observed since sucrose already inhibits SnRK1 activity. Consistent with negative regulation of SnRK1 activity by T6P (Zhang et al., 2009), the *tps1* mutant, deficient in the major T6P synthase TPS1 (Eastmond et al., 2002; Van Dijken et al., 2004), shows significantly reduced anthocyanin accumulation and *MYB75* and *DFR* expression in sucrose-rich medium (Figure 2C,D). Moreover, the recently described *tps1* suppressor mutants with additional mutations in the SnRK1α1 catalytic site (160-1: G178R, 199-6: R259Q, 232-2: G162D) and concomitant restoration of embryogenesis and transition to flowering (Zacharaki et al., 2022) showed a fully restored sucrose-induced anthocyanin biosynthesis and *MYB75* and *DFR* expression (Figure 2E,F). Together, our results confirm that SnRK1 is an important negative regulator of sucrose-induced anthocyanin biosynthesis, at least in part by transcriptional repression of *MYB75*, a key factor in controlling expression of *DFR*, an essential enzyme for anthocyanin biosynthesis.

**Figure 2.**
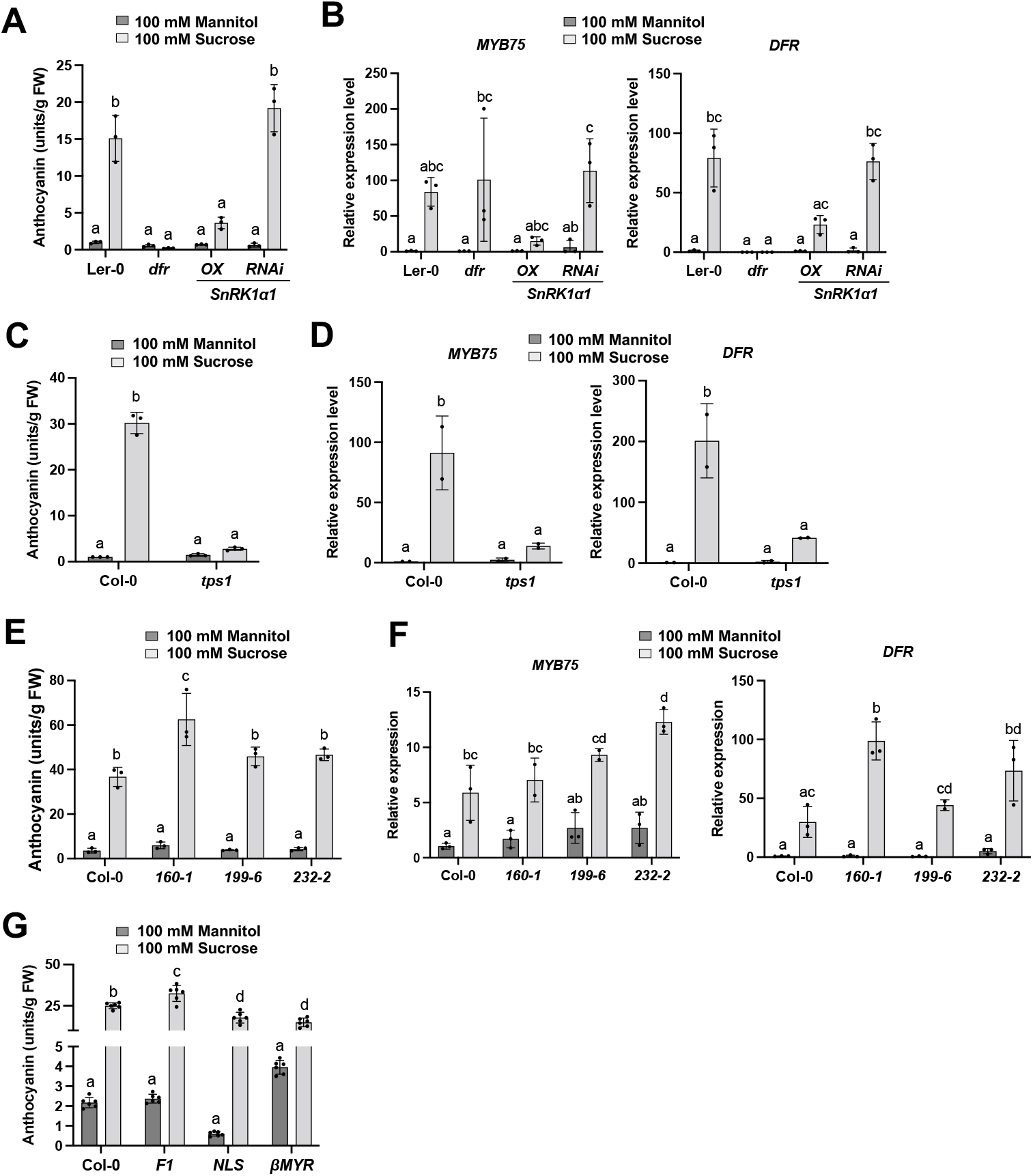
SnRK1 represses sucrose-induced anthocyanin biosynthesis. (A) Relative anthocyanin levels of 7-day-old Ler-0 wildtype, dfr, SnRK1α1 OX and SnRK1α1 RNAi seedlings grown in ½ MS medium supplemented with 100 mM mannitol or 100 mM sucrose. Values are averages with SD, n = 3 biological repeats. Two-way ANOVA statistical analysis was performed in GraphPad Prism v9, letters represent statistically significant differences, p<0.0001. FW, fresh weight. (B) qRT-PCR analysis of MYB75 and DFR gene expression levels in 7-day-old Ler-0 wildtype, dfr, SnRK1α1 OX and SnRK1α1 RNAi seedlings grown in ½ MS medium supplemented with 100 mM mannitol or 100 mM sucrose. Values are averages with SD, n = 3 biological repeats. Two-way ANOVA statistical analysis was performed in GraphPad Prism v9, letters represent statistically significant differences, MYB75: p<0.05 – DFR: p<0.0001. (C) Relative anthocyanin levels of 7-day-old Col-0 wildtype and tps1 seedlings grown in ½ MS medium supplemented with 100 mM mannitol or 100 mM sucrose. Values are averages with SD, n = 3 biological repeats. Two-way ANOVA statistical analysis was performed in GraphPad Prism v9, letters represent statistically significant differences, p<0.0001. FW, fresh weight. (D) qRT-PCR analysis of MYB75 and DFR gene expression levels in 7-day-old Col-0 wildtype and tps1 seedlings grown in ½ MS medium supplemented with 100 mM mannitol or 100 mM sucrose. Values are averages with SD, n = 2 biological repeats. Two-way ANOVA statistical analysis was performed in GraphPad Prism v9, letters represent statistically significant differences, p<0.025. (E) Relative anthocyanin levels in wildtype Col-0 and three tps1-2 GVG:TPS1 (tps1) suppressor mutants seedlings (with mutations in SnRK1α1) grown for 7 days in ½ MS medium supplemented with 100 mM mannitol or 100 mM sucrose. 160-1: SnRK1α1 G178R, 199-6: R259Q, 232-2: G162D. Values are averages with SD, n = 3 biological repeats. Two-way ANOVA statistical analysis was performed in GraphPad Prism v9, letters represent statistically significant differences, p<0.016. FW, fresh weight. (F) qRT-PCR analysis of MYB75 and DFR gene expression levels in Col-0 wildtype and three tps1-2 GVG:TPS1 (tps1) suppressor mutants seedlings (160-1, 199-6 and 232-2) grown in ½ MS medium supplemented with 100 mM mannitol or 100 mM sucrose. Values are averages with SD, n = 3 biological repeats. Two-way ANOVA statistical analysis was performed in GraphPad Prism v9, letters represent statistically significant differences, p<0.05. (G) Relative anthocyanin levels of 7-day-old Col-0 wildtype and SnRK1α1/SnRK1α1 SnRK1α2/ SnRK1α2 double mutant seedlings complemented with wildtype SnRK1α1 (F1), NLS-SnRK1α1 (NLS), and βMYR-SnRK1α1 (βMYR) seedlings grown in ½ MS medium supplemented with 100 mM mannitol or 100 mM sucrose. Values are averages with SD, n = 6 biological repeats. Twoway ANOVA statistical analysis was performed in GraphPad Prism v9, letters represent statistically significant differences, p<0.0001. FW, fresh weight.

Previous work identified an important regulatory role for subcellular localization in SnRK1 signaling, with SnRK1α translocating to the nucleus upon metabolic stress to activate target gene induction. However, target gene (including *MYB75*) repression, does not require nuclear localization (Ramon et al., 2019). We quantified anthocyanin levels in WT and transgenic seedlings with either increased nuclear (NLS-SnRK1α1) or increased cytoplasmic (βMYR-SnRK1α1) localization of SnRK1α1 (Ramon et al., 2019). Anthocyanin levels did not differ significantly between lines on sucrose medium, which is consistent with the repression of SnRK1 activity in sugar rich conditions. In osmotic control conditions (100 mM mannitol), anthocyanin levels were significantly lower in the NLS seedlings and significantly higher in βMYR seedlings (Figure 2G). Since target gene repression does not require nuclear translocation, the latter observation points to a non-transcriptional inhibitory effect of nuclear SnRK1α1. In addition, the short-term transcriptional effects observed in seedlings (Figure S2A) are consistent with a more complex regulation. Interestingly, we previously observed that after flowering (during senescence) NLS-SnRK1α1 lines accumulate significantly less anthocyanins (Figure S2B).

### Post-translational repression of MBW complex activity by SnRK1

To explore possible post-translational regulation of nuclear MBW complex activity by SnRK1, we introduced a *35S::SnRK1α1* construct in the *pap1-D* mutant. *SnRK1α1 OX* appears to suppress the increased anthocyanin accumulation (Figure S2C). However, we also found that *MYB75* expression was also significantly repressed (Figure S2D), indicating that *MYB75* expression (transcript level) is indeed still subject to metabolic status and SnRK1 regulation in the *pap1-D* mutant. Unfortunately, while we confirmed BASTA (*pap1-D*) and kanamycin (*35S::SnRK1α1* construct) resistance and the presence of the *35S::SnRK1α1* construct (by PCR and sequencing) in 4 independent lines, we were not able to detect increased *SnRK1α1* expression levels. This suggests silencing and needs to be resolved. We therefore turned to cellular assays that have previously proven very helpful in elucidating SnRK1 signaling mechanisms (Ramon et al., 2019). We transiently expressed the MYB75, bHLH2 and TTG1 MBW complex subunits in *Arabidopsis* leaf mesophyll protoplasts, eliminating transcriptional regulation by using a constitutive 35S promoter. A 1000 bp *DFR* promoter driving firefly luciferase (LUC) expression (Pr*DFRA::LUC*) was used as a reporter for MBW complex activity. While transient overexpression of MYB75 led to a significant increase in Pr*DFRA::LUC* activity, overexpression of bHLH2 and TTG1 did not (Figure 3A,B). However, a strong synergistic effect on Pr*DFRA::LUC* activity was observed upon co-expressing MYB75 and bHLH2, with a further increase upon co-expression of TTG1 (Figure 3A,B). This confirmed the formation of a functional MBW complex in this setup. When we looked at the effect on endogenous *DFR* promoter activity using qRT-PCR, *MYB75* overexpression by itself did not significantly trigger expression, but a significant synergistic effect was seen upon bHLH2 co-expression, that was not further increased with additional *TTG1* overexpression (Figure 3C). However, the strong effect of TTG1 co-expression in *ttg1* KO protoplasts confirmed the essential role of this protein for optimal activity of the MYB and bHLH TFs (Figure 3C).

**Figure 3.**
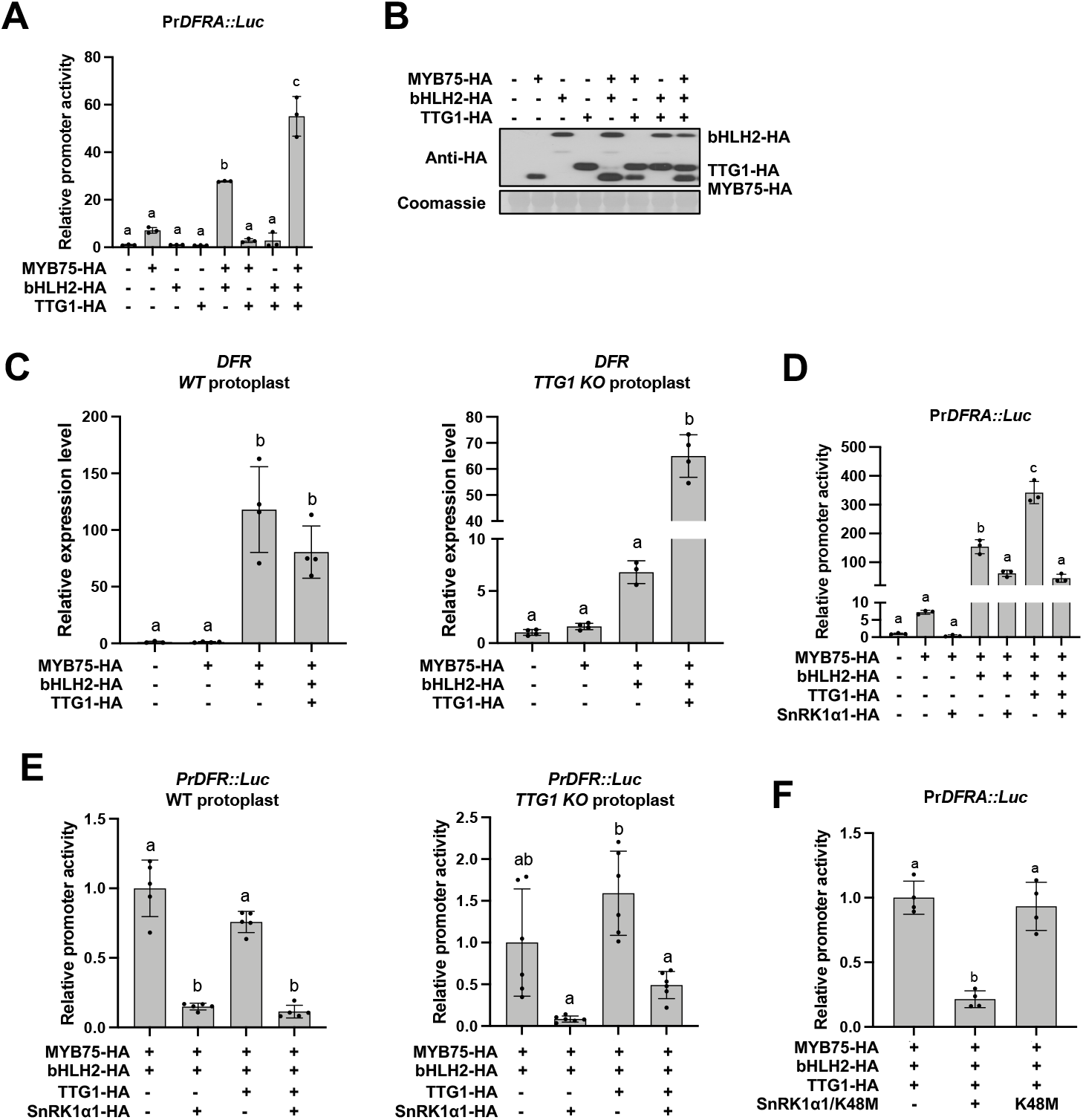
SnRK1α1 inhibits MBW complex activation of the DFR promoter in leaf mesophyll protoplasts. (A) DFR promoter activity in Arabidopsis leaf mesophyll protoplasts upon transient coexpression of MYB75, bHLH2 and TTG1. (B) Expression assessment of HA-tagged proteins by immunoblot analysis with anti-HA antibodies, using RBCS staining with Coomassie Brilliant Blue R-250 as a protein loading control. (C) qRT-PCR analysis of DFR gene expression in Col-0 wildtype and ttg1 mutant leaf mesophyll protoplasts upon transient of MYB75, bHLH2 and TTG1. (D) DFR promoter activity in Col-0 wildtype or ttg1 KO Arabidopsis leaf mesophyll protoplasts upon transient co-expression of MYB75, bHLH2 and TTG1 with SnRK1α1. (E) DFR promoter activity in Arabidopsis leaf mesophyll protoplasts upon transient coexpression of MYB75, bHLH2 and TTG1 with SnRK1α1. (F) DFR promoter activity in Arabidopsis leaf mesophyll protoplasts upon transient coexpression of the full MBW complex with wildtype SnRK1α1 or the kinase dead SnRK1α1 K48M mutant protein. Relative and normalized promoter activity values are averages with SD, n = 3 (A), n = 4 (C), n = 3 (D), n = 5 (E), n = 4 (F) biological repeats (independent protoplast transfections). One-way ANOVA statistical analysis was performed in GraphPad Prism v9, letters represent statistically significant differences, p<0.005.

*MYB75* expression was first identified as a critical component in sucrose-specific induction of anthocyanin biosynthesis in a QTL analysis (Teng et al., 2005). The preceding analysis of 43 *Arabidopsis* accessions revealed a considerable natural variation in sucrose-induced anthocyanin accumulation. Interestingly, the Cvi ecotype is almost non-responsive; sucrose can still induce *MYB75* expression but does not trigger anthocyanin accumulation, suggesting that the Cvi MYB75 protein is not functional, likely due to the two unique SNPs resulting in P37H and K160N mutations compared to the Col and Ler ecotypes (Teng et al., 2005). We therefore separately tested the effect of the P37H and K160N mutations on the MYB75 protein’s ability to activate the Pr*DFRA::LUC* reporter in leaf mesophyll protoplasts. Mutation of P37 completely abolished transcription activity of MYB75, while K160N had no effect (Figure S3A). Online assessment of the protein predicted features of MYB75 protein upon P37H mutation suggested disruption of the alpha helix and a change in protein binding properties (Yachdav et al., 2014). P37 is localized in the R2 motif of the MYB75 MYB domain. Based on the crystal structure of the well-characterized R2R3 MYB protein MYB domain (WER), the MYB domain of Col and Cvi MYB75 (59.62% identity with WER) was modeled using Swiss-Model (Waterhouse et al., 2018). In contrast to the above mentioned online prediction, structural disturbance of the alfa helix upon Pro37 mutation to His is limited (Cvi model) (Figure S3B). This may suggest that the lack of *DFR* promoter activity is caused with changes in protein interaction properties rather than having a structural origin.

Interestingly, SnRK1α1 co-expression significantly repressed the activating effect of MYB75, MYB75 and bHLH2, and the full MBW complex on the Pr*DFRA::LUC* reporter, both in Col-0 WT (Figure 3D,E) and in *ttg1* KO protoplasts (Figure 3E). This effect is dependent on SnRK1α1 kinase activity. Expression of the kinase dead SnRK1α1 K48M mutant protein had no effect (Figure 3F), indicating that the repressive effect of SnRK1 is mediated by phosphorylation of one or more MBW complex subunits or upstream regulators. While we focused further research on the *DFR* promoter, overexpression of the MBW complex also more generally activated endogenous expression of the LBGs, including *F3H, LDOX* and *UF3GT*, and not the flavonol biosynthetic gene *FLS* (Figure 4A). In all cases, this activation was also subject to the kinase activity-dependent repressive effect of SnRK1α1 co-expression (Figure 4B). These results indicate that transient overexpression of the MBW complex in leaf mesophyll protoplasts can be used as an experimental system to explore the molecular mechanisms involved in its post-translational regulation by SnRK1.

**Figure 4.**
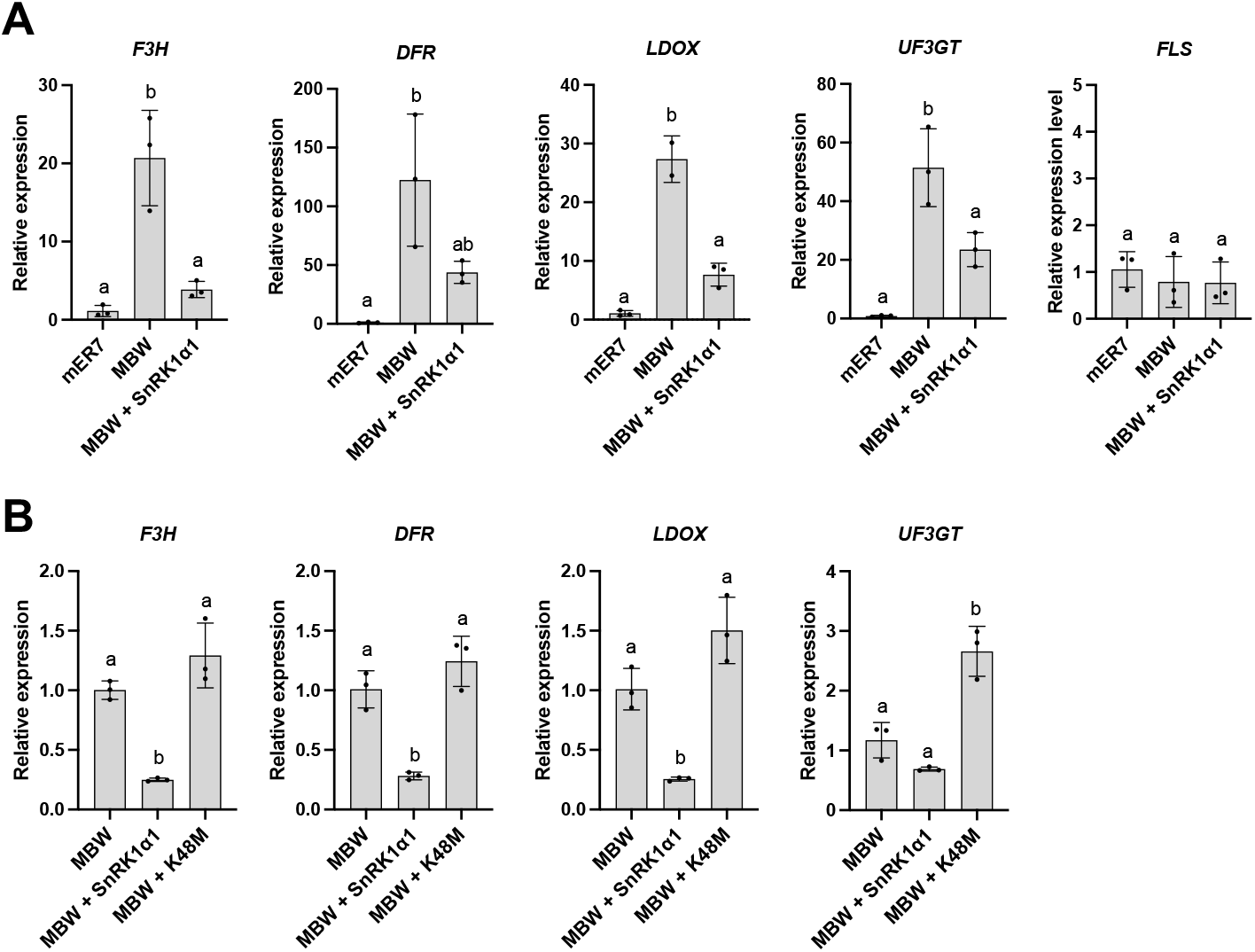
SnRK1α1 inhibits MBW complex activation of LBGs in leaf mesophyll protoplasts. (A) qRT-PCR analysis of gene expression levels of late biosynthetic genes (F3H, DFR, LDOX, UF3GT and FLS) in Arabidopsis leaf mesophyll protoplasts upon transient co-expression of the MBW complex without or with SnRK1α1. (B) qRT-PCR analysis of gene expression levels of late flavonoid biosynthesis enzymes (F3H, DFR, LDOX, UF3GT and FLS) in Arabidopsis leaf mesophyll protoplasts upon transient coexpression of the MBW complex with wildtype SnRK1α1 or the kinase dead SnRK1α1 K48M mutant protein. Values are averages with SD, n = 3 biological repeats. One-way ANOVA statistical analysis was performed in GraphPad Prism v9, letters represent statistically significant differences, p<0.01.

### SnRK1 triggers MBW complex dissociation and MYB75 protein degradation

We first studied the effect of SnRK1 on MBW subunit interactions using coimmunoprecipitation. Transiently expressed FLAG-tagged MYB75, bHLH2 and TTG1 proteins were pulled down (using anti-FLAG beads) and co-immunoprecipitating co-expressed HA-tagged MBW complex subunits were visualized by immunoblotting using anti-HA antibodies (Figure 5A). We found clear co-immunoprecipitation of the respective other two MBW complex proteins, confirming effective trimeric complex formation in this setup. Coexpression of HA-tagged SnRK1α1 reduced the amounts of co-immunoprecipitating proteins. This effect was not observed with co-expression of the catalytically dead SnRK1α1 K48M mutant protein, indicating complex dissociation by SnRK1 activity.

**Figure 5.**
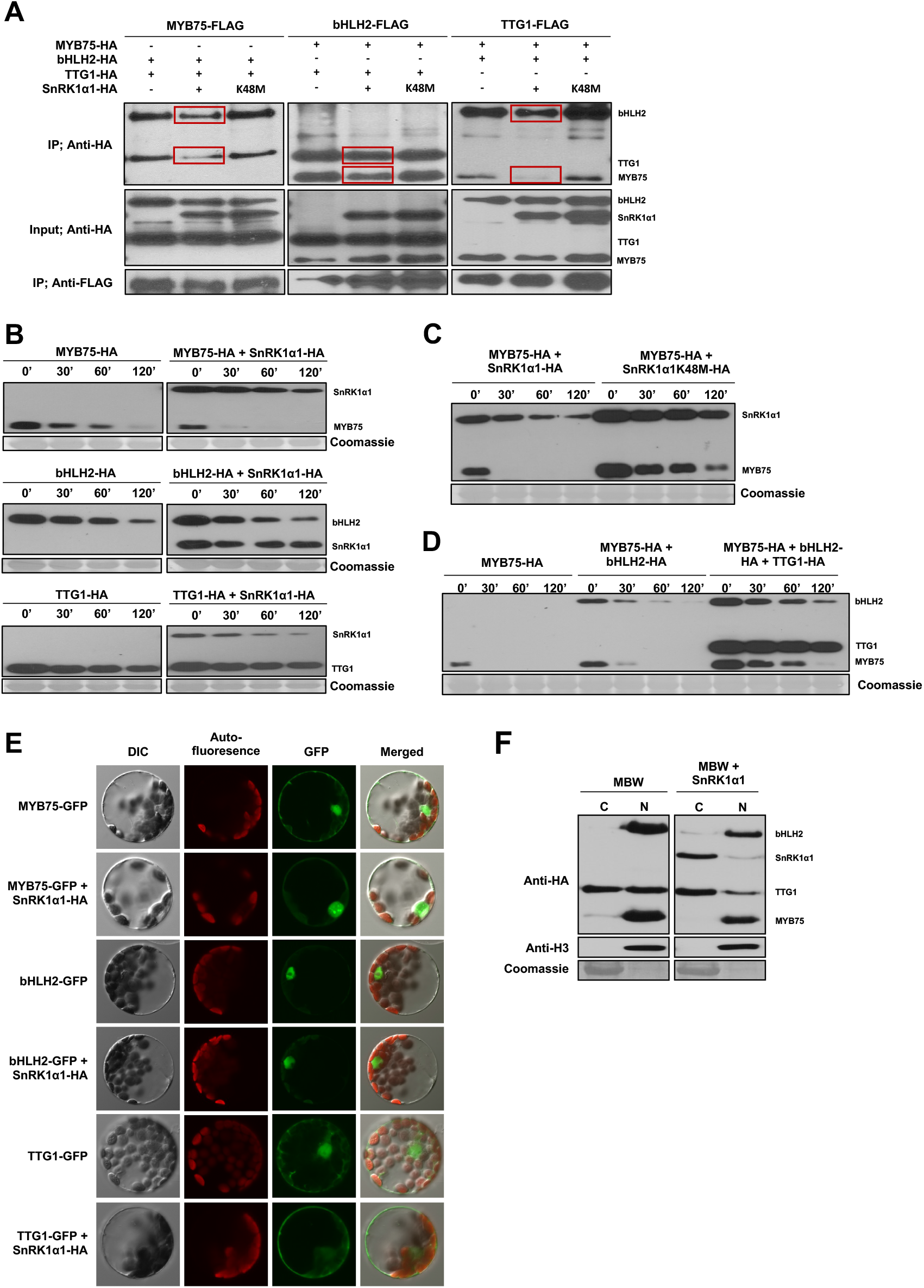
SnRK1α1-induced MBW complex dissociation, MYB75 protein degradation and TTG1 protein translocation. (A) Co-immunoprecipitation of HA-tagged MBW subunits with FLAG-tagged MBW without and with co-expression of wildtype or kinase dead K48M mutant SnRK1α1. Protein input and IP were visualized by immunoblot analysis using anti-HA and anti-FLAG antibodies, as indicated. (B) MBW subunit stability without and with SnRK1α1 co-expression in leaf mesophyll protoplasts. HA-taggedprotein levels are analysed 0, 30, 60 and 120 minutes after addition of 10 μM cycloheximide (CHX) by immunoblotting with anti-HA antibodies. MG132 (10μM) is used as a proteasome inhibition control. RBCS staining with Coomassie Brilliant Blue R-250 serves as protein loading controls. (C) and (D) MYB75 protein stability without and with co-expression of wildtype or kinase dead K48M mutant SnRK1α1 (C) or the other MBW complex subunits (D) in leaf mesophyll protoplasts. HA-tagged protein levels are analyzed 0, 30, 60 and 120 minutes after addition of 10 μM cycloheximide (CHX) by immunoblotting with anti-HA antibodies. MG132 (10μM) is used as a proteasome inhibition control. RBCS staining with Coomassie Brilliant Blue R-250 serves as protein loading controls. (E) Subcellular localization of MYB75-GFP, bHLH2-GFP and TTG1-GFP in leaf mesophyll protoplasts without and with SnRK1α1 co-expression, analyzed by fluorescence microscopy 16 h after transfection. DIC, differential interference contrast image. (F) Immunoblot analysis of nuclear (N) and cytoplasmic (C) fractions of HA-tagged MBW subunit proteins without and with SnRK1α1 co-expression in leaf mesophyll protoplast using anti-HA antibodies. Anti-Histone H3 antibodies and RBCS staining with Coomassie Brilliant Blue R-250 serve as controls for purity of the nuclear and cytoplasmic fractions, respectively. Ten percent of the cytoplasmic fractions and the complete nuclear fractions of samples were used for analysis.

MYB75 protein stability is known to be regulated by proteasomal degradation in response to light (Mair et al., 2013) and also bHLH2/EGL3 is targeted for degradation (Patra et al., 2013). Transient expression of the different MBW subunits in leaf mesophyll protoplasts and immunoblot analysis of protein levels after cycloheximide (CHX) treatment (blocking new protein synthesis) confirmed that MYB75 and bHLH2, but not TTG1, are unstable proteins. Interestingly, co-expression of SnRK1α1 significantly accelerated MYB75 protein degradation (Figure 5B). No obvious effects of SnRK1α1 on bHLH2 or TTG1 protein stability were observed. The destabilizing effect of SnRK1α1 on MYB75 also depends on its phosphorylation capacity as MYB75 stability was not affected upon co-expression of the mutant SnRK1α1 K48M protein (Figure 5C). We also saw an increase in MYB75 stability upon co-expression of the other MBW complex subunits (Figure 5D), suggesting that MYB75 is degraded upon complex dissociation. While the MYB and bHLH TFs are typically localized in the nucleus, TTG1 is known to shuttle between nucleus and cytosol. Analyzing the subcellular localization of the eGFP-tagged MBW subunits using confocal microscopy, we observed a shift of TTG1 from a predominantly nuclear to a more cytosolic localization upon co-expression of SnRK1α1 (Figure 5E). Immunoblotting of nuclear and cytosolic fractions of transfected leaf mesophyll protoplasts confirmed reduced amounts of TTG1 in the nuclear fraction when SnRK1α1 was co-expressed. This translocation further corroborates the dissociation of the MBW complex by SnRK1 activity. The localization of MYB75 and bHLH2 appeared unaffected (Figure 5E,F).

### SnRK1 triggers MYB75 release from *DFR* promoter chromatin

We then turned our attention to the *DFR* promoter, first progressively truncating the sequence to identify putative TF binding sites. The sequence between −350 and −300 bp relative to the start codon appeared essential for activation by the MBW complex (Figure 6A; Figure S4A). Interestingly, this 50 bp sequence contains a G-box-like CACGTC sequence as a candidate bHLH TF binding site (Figure 6B). Previously, a motif analysis of 8 promoters (including that of *DFRA*) transactivated in a leaf infiltration assays identified the conserved (C/T)CNCCAC(A/G)(A/T)(G/T) or (C/T)(A/C)NCCACN(G/T)(G/T) motif with core CCAC sequence as a cis-regulatory element required for activation by MYB75 (Dare et al., 2008). Five of the promoters contained a CACGTG G-box site adjacent to the CCAC sequence suggesting that it may not correspond to the binding site of MYB75, but that of the associated bHLH TF (Dare et al., 2008). In the 350 bp *DFR* promoter sequence, we found two such CCAC motifs associated with a perfect CACGTG G-box and one with the CACGTC G-box-like sequence (Figure S4B). MYB TF binding sites are less well characterized. However, the petal-specific flavonoid biosynthesis regulator MYB305 from Snapdragon (*Antirrhinum majus*) was shown to bind a conserved (A/C)ACC(A/T)A(A/C)C sequence (Sablowski et al., 1994). We found a consistent CACCAAAC sequence right next to the putative bHLH binding site in the 50 bp stretch between −350 and −300 bp relative to the start codon. Interestingly, although *PAL2* is an EBG, activation of the *PAL2* promoter by MYB305 in tobacco protoplasts also required a nearby G-box-like CACGTC element (Sablowski et al., 1994). We mutated the putative bHLH (G-box-like sequence; 5’-CACGTC-3’) and MYB (MYB-core element; 5’-CACCAAAC-3’) binding sites and found that both mutations reduced promoter activity upon co-expression of the MBW complex (Figure 6C). Mutation of the CACGTC sequence had the most significant effect. Double mutation reduced promoter activity even further, to the same extend as truncation of the promoter to 300 bp. This indicates that the mutated elements are indeed required for proper activation of *DFR* expression and likely function as the MBW recognition binding site elements.

**Figure 6.**
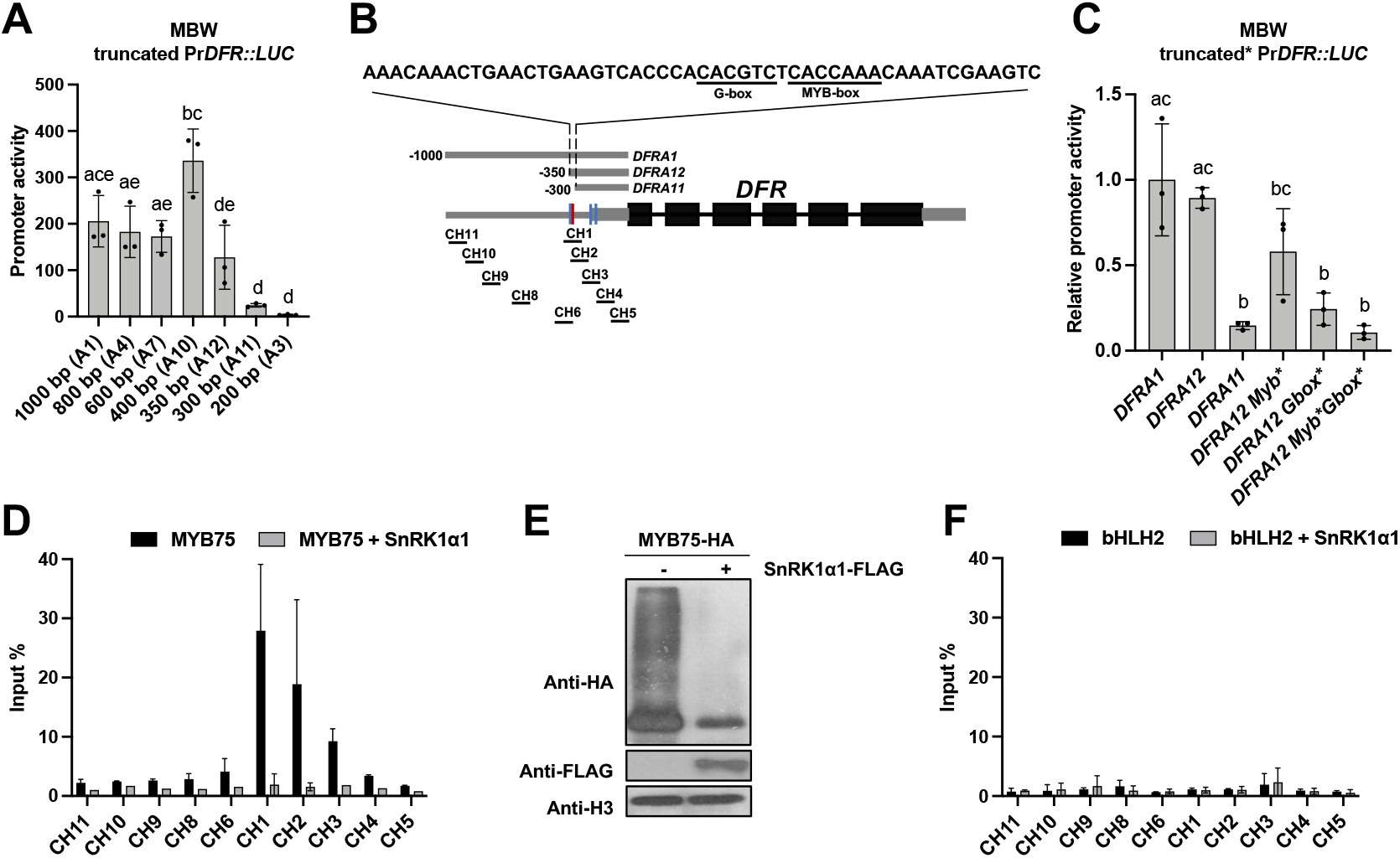
MYB75 binding to and SnRK1α1-induced dissociation from the DFR promoter. (A) DFR promoter activity with transient MBW complex expression in Arabidopsis leaf mesophyll protoplasts upon progressive sequence truncation. Values are averages with SD, n = 3 biological repeats (independent protoplast transfections). One-way ANOVA statistical analysis was performed in GraphPad Prism v9, letters represent statistically significant differences, p<0.05. (B) Schematic representation of the DFR promoter, the key 50 bp sequence and motifs identified, and the position of the PCR amplicons used for ChIP analysis (CH1-11). (C) Truncated and mutated DFR promoter activity upon overexpression of the MBW complex in Arabidopsis leaf mesophyll protoplasts. The G-box-like (CACGTC) and MYB core (CACCAAAC) elements in the 350 bp promoter (DFRA12) were mutated. Values are averages with SD, n = 3 biological repeats (independent protoplast transfections). One-way ANOVA statistical analysis was performed in GraphPad Prism v9, letters represent statistically significant differences, p<0.01. (D) PCR analysis of DFR promoter sequences (input %) co-precipitating with immunoprecipitated HA-tagged MYB75 (ChIP) with and without SnRK1α1 co-expression. (E) Levels of immunoprecipitated MYB75 protein without and with SnRK1α1 co-expression. (F) PCR analysis of DFR promoter sequences (input %) co-precipitating with immunoprecipitated HA-tagged bHLH2 (ChIP) with and without SnRK1 co-expression.

We then performed a chromatin immunoprecipitation (ChIP) analysis using FLAG-tagged MYB75 overexpression in leaf mesophyll cells. This confirmed that the sequence containing the putative G- and MYB-box is important for MYB75 binding (Figure 6D). Co-expression of SnRK1α1 completely abolished ChIP indicating MYB75 TF release from the promoter chromatin, although reduced enrichment is also partially due to reduced MYB75 protein stability (Figure 6E). ChIP with bHLH2-FLAG did not yield any enrichment (Figure 6F), suggesting that MYB75 binds first to then recruit the other complex members to the promoter.

### SnRK1α1 directly interacts with and phosphorylates the MBW complex

Our results indicate that SnRK1 kinase activity is required to inhibit the MBW complex. We first assessed direct interaction of SnRK1α1 with the MBW complex subunits via coimmunoprecipitation. Transiently expressed FLAG-tagged SnRK1α1 was pulled down using anti-FLAG beads and co-immunoprecipitating HA-tagged MBW complex subunits were visualized by immunoblotting using anti-HA antibodies (Figure 7A). This suggests interaction of SnRK1α1 with each of the three MBW subunits, MYB75, bHLH2 and TTG1. To determine whether and which MBW subunits are also phosphorylated by SnRK1 we first performed a Phos-tag™ SDS-PAGE mobility shift assay with all three MBW subunit without and with coexpression of SnRK1α1 or the SnRK1α1 K48M kinase dead protein. Interestingly, MYB75 and TTG1 produced respectively one and two SnRK1α1-dependent phosphorylation bands (Figure 7B).

**Figure 7.**
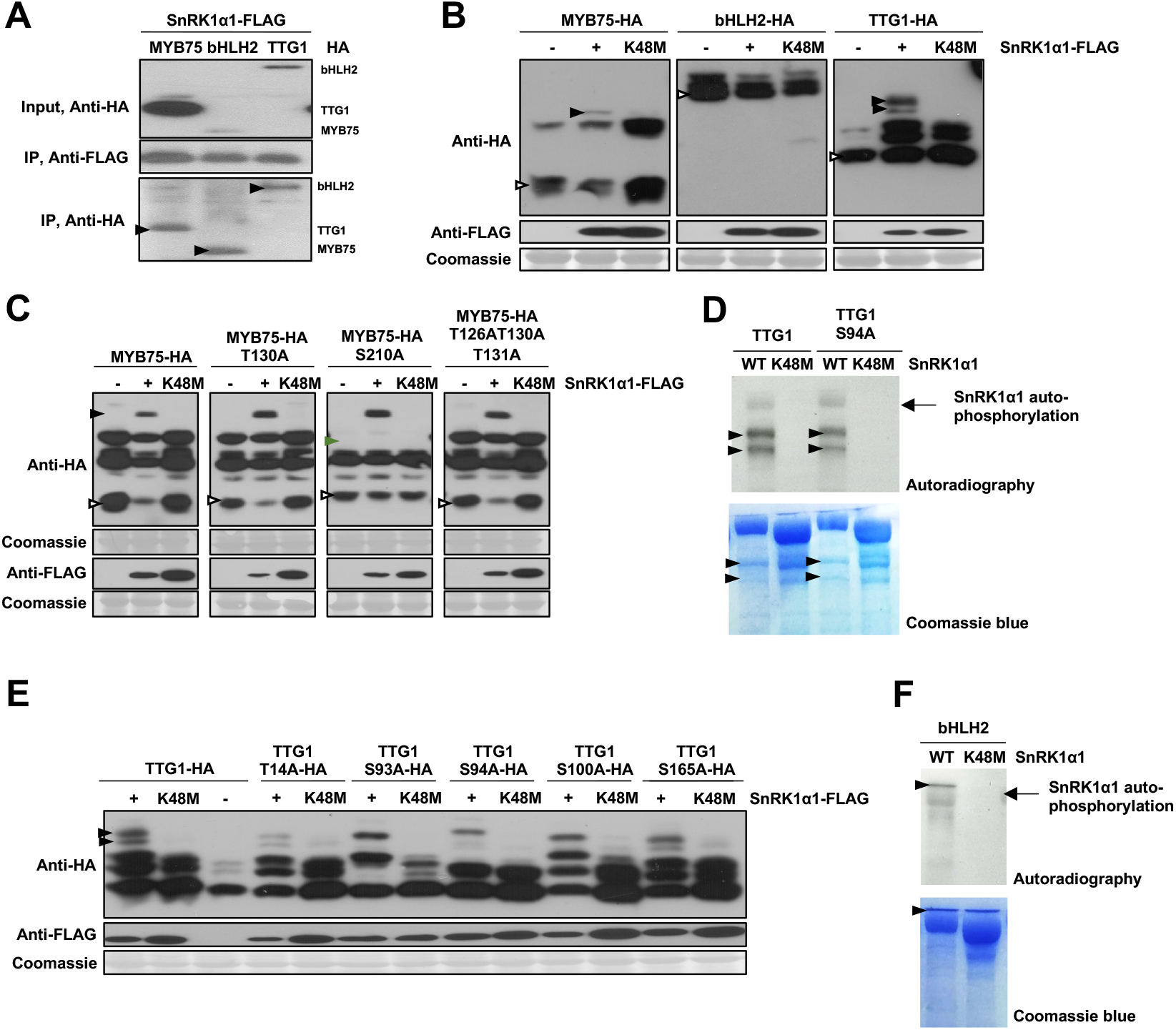
SnRK1α1 interacts with and phosphorylates all three MBW complex subunits, MYB75, bHLH and TTG1. (A) The MBW subunits MYB75, bHLH2 and TTG1 all interact with SnRK1α1. Coimmunoprecipitation of transiently co-expressed HA-tagged MYB75, bHLH2 and TTG1 with SnRK1α1-FLAG in Arabidopsis leaf mesophyll protoplasts. Protein input and IP were visualized by immunoblot analysis using anti-HA and anti-FLAG antibodies, as indicated. (B) Phos-tag acrylamide-based mobility shift assay of MYB75-HA, bHLH2-HA and TTG1-HA proteins expressed in leaf mesophyll protoplasts without and with co-expression of wildtype or kinase dead K48M mutant SnRK1α1. Black arrows indicate SnRK1α1-specific phosphorylated protein bands, white arrows indicate non-phosphorylated proteins. Immunoblot analysis was performed using anti-HA and anti-FLAG antibodies and RBCS staining with Coomassie Brilliant Blue R-250 as a protein loading control. (C) Phos-tag acrylamide-based mobility shift assay of MYB75-HA proteins with mutated putative SnRK1-phosphorylated residues, transiently expressed in leaf mesophyll protoplasts without and with co-expression of wildtype or kinase dead K48M mutant SnRK1α1. Black arrows indicate SnRK1α1-specific phosphorylated protein bands. Green arrow indicates loss of protein phosphorylation band, white arrows indicate non-phosphorylated proteins. Immunoblot analysis was performed using anti-HA and anti-FLAG antibodies and RBCS staining with Coomassie Brilliant Blue R-250 as a protein loading control. (D) An in vitro kinase assay with His6-MBP-tagged wildtype or mutated (S94A) TTG1 and SnRK1α1 and SnRK1α1 K48M proteins. The proteins were separated after the kinase reaction via SDS-PAGE, analyzed by autoradiography and stained with Coomassie Brilliant Blue R-250. (E) Phos-tag acrylamide-based mobility shift assay of TTG1-HA proteins with mutated putative SnRK1-phosphorylated residues, transiently expressed in leaf mesophyll protoplasts without and with co-expression of wildtype or kinase dead K48M mutant SnRK1α1. Black arrows indicate SnRK1α1-specific phosphorylated protein bands. Green arrow indicates loss of protein phosphorylation band. Immunoblot analysis was performed using anti-HA and anti-FLAG antibodies and RBCS staining with Coomassie Brilliant Blue R-250 as a protein loading control. (F) An in vitro kinase assay with His6-MBP-tagged bHLH2 and SnRK1α1 and SnRK1α1 K48M proteins. The proteins were separated after the kinase reaction via SDS-PAGE, analyzed by autoradiography and stained with Coomassie Brilliant Blue R-250.

Based on the 10 amino acid (M, L, V, I, F) XXXX S(T) XXX (M, L, V, I, F) consensus recognition motif, MYB75 contains two putative SnRK1α1 phosphorylation sites, T130 and S210. This motif starts and ends with a bulky hydrophobic residue (M, L, V, I, or F) at positions P-5 and P+4 relative to the phosphorylated serine or threonine, preferably with a basic residue (R>K>H) at position P-3 or P-4. Li et al (2016) previously identified T130 as a phosphorylated residue without identifying the responsible kinase. In addition, MYB75 was found to be phosphorylated at T126 and T131 by MAPK4, increasing MYB75 stability (Li al., 2016). Mutating the MYB75 (putative) phosphorylation sites T126, T130 and T131 to an alanine (A) residue, however, did not alter the phosphorylation pattern in the Phos-tag™ SDS-PAGE mobility shift assay (Figure 7C). Mutation of S210 into an alanine residue, on the other hand, resulted in loss of one phosphorylation band both without and with SnRK1α1 or SnRK1α1 K48M co-expression. This result identifies S210 as a phosphorylated MYB75 residue, however, its phosphorylation does not seem to be SnRK1α1 dependent. Purification of recombinant His6-MBP-tagged MYB75 did not yield sufficient protein for *in vitro* kinase and mass spectrometry analyses.

The TTG1 protein sequence only contains a single perfect consensus SnRK1 recognition motif, around S94. An *in vitro* kinase assay with His6-MBP-tagged TTG1 and SnRK1α1 and SnRK1α1 K48M proteins confirmed that TTG1 is phosphorylated by SnRK1 and that S94A mutation did not abolish phosphorylation (Figure7D). Subsequent mass Spectrometry (MS) analysis identified five SnRK1α1 phosphorylation sites with high probability: T14, S93, S94, S100 and S165 (Table S3). We also mutated the other 4 residues into alanine. The altered phosphorylation patterns of the S93A, S94A and S100A mutant TTG1 proteins in a Phos-tag™ SDS-PAGE mobility shift assay confirmed that these residues are indeed SnRK1 phosphorylation sites (Figure 7E).

bHLH2 did not produce a distinct pattern in the mobility shift assay with SnRK1α1 coexpression, but negative results are not conclusive. We therefore also performed a kinase assay with His6-MBP-tagged bHLH2 and SnRK1α1 and SnRK1α1 K48M proteins and this indicates that also bHLH2 is a direct SnRK1 phosphorylation target (Figure 7F).

Cellular assays were then used to further investigate whether MYB75 S210 or TTG1 S93A S94A S100A mutation affects MBW complex activity. MBW complexes containing a MYB75 S210A mutant protein showed reduced activation of the Pr*DFR::LUC* reporter, but SnRK1α1 coexpression still repressed the complex to the same basal activity (Figure S5A). No effect could be observed for the triple TTG1 mutant (Figure S5B), probably due to presence of additional SnRK1 phosphorylation sites. Relaxing the consensus recognition motif indeed identifies additional putative SnRK1α1 phosphorylation sites, such as S60, S126, T158, S193, S197, S242 and S252.

In conclusion, SnRK1α1 directly interacts with all three MBW complex proteins on multiple residues. Which combinations are essential for regulation remains to be determined.

### SnRK1 induces expression of the negative MBW complex regulator MYBL2

Several negative regulators have been identified in *Arabidopsis* which further fine-tune MBW complex activity. The single repeat R3-MYB protein MYBL2 was shown to competitively bind with the bHLH proteins GL3, EGL3 and bHLH42/TT8 in yeast (Sawa, 2002; Zimmermann et al., 2004), preventing the formation of a functional MBW complex (Dubos et al., 2008; Matsui et al., 2008). We confirmed this in our cellular assay. Co-expression of MYBL2 repressed Pr*DFR::LUC* reporter activation by the MBW complex (Figure 8A). Interestingly, MYBL2 appears to be a (direct or indirect) transcriptional target of SnRK1 as its expression was significantly upregulated in leaf mesophyll protoplasts overexpressing SnRK1α1 (Figure 8B). This response is also confirmed in a seedling starvation assay. Removal of sucrose from 7-day-old sucrose-grown Col-0 seedlings triggered an increase in *MYBL2* expression, while repressing *MYB75* and *DFR* expression (Figure 8C). This provides yet another means for SnRK1 to inhibit MBW complex activity and downregulate anthocyanin biosynthesis.

**Figure 8.**
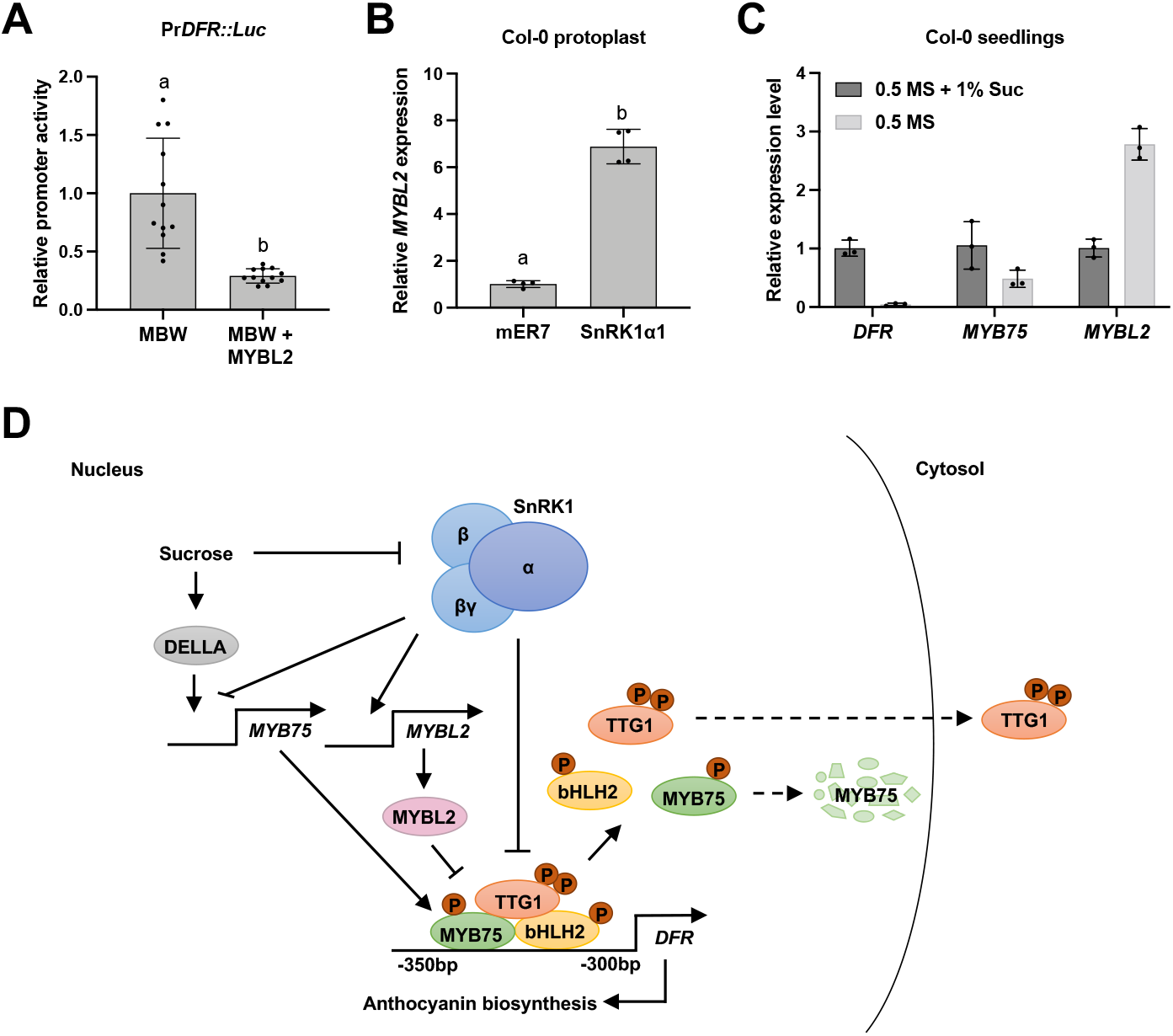
Integration of MYBL2 in a model for anthocyanin regulation by SnRK1. (A) DFR promoter activity in leaf mesophyll protoplasts upon transiently expressing MBW complex without and with co-expression of MYBL2. Values are averages with SD, n = 12 biological repeats (independent protoplast transfections). Unpaired t-test analysis was performed in GraphPad Prism v9, letters represent statistically significant differences, p<0.0001. (B) qRT-PCR analysis of endogenous MYBL2 expression in leaf mesophyll protoplasts without and with transient SnRK1α1 overexpression. Values are averages with SD, n = 4 biological repeats. Unpaired t-test analysis was performed in GraphPad Prism v9, letters represent statistically significant differences, p<0.0001. (C) qRT-PCR analysis of endogenous DFR, MYB75 and MYBL2 expression in non-sugar-starved and sugar-starved 7-day-old Col-0 seedlings. Values are averages with SD, n = 3 biological repeats (15 pooled seedlings each). (D) Model for multi-level anthocyanin biosynthesis regulation by SnRK1. Transcriptional regulation of the MBW complex involves repression of MYB75 expression, possibly independent of the DELLA proteins, and induction of the negative regulator MYBL2. Post-translational regulation involves direct interaction with and phosphorylation of MYB75, TTG1 and bHLH2, associated with complex dissociation, accelerated MYB75 degradation, and translocation of TTG1 to the cytosol.

## Discussion

The physiological relevance of sucrose-induced anthocyanin accumulation is most obvious for sugar-rich ripe fruits, attracting animals for seed dispersal, and vulnerable proliferating vegetative sink tissues. In stressful conditions that still allow photosynthesis and sucrose biosynthesis but disable photosynthate utilization or export, such as cold stress (in which sucrose also plays an important role as primary osmolyte), and under high light intensities and UV radiation, anthocyanin accumulation protects against concomitant oxidative stress and acts as a sunscreen. But while some stress conditions are associated with sugar accumulation, others result in C and energy depletion because of direct or indirect effects on photosynthesis, respiration, or C allocation. The energy saving mode triggered by SnRK1 activation in such metabolic stress conditions results in the suppression of energy-consuming anabolic and growth processes, redirecting C flux to more essential functions ensuring survival (Broeckx et al., 2016). This is clearly illustrated by the inverse correlation between SnRK1 activity and expensive anthocyanin biosynthesis. *Arabidopsis SnRK1α1/KIN10* overexpression and estradiol-induced *SnRK1α1/kin10 SnRK1α2/kin11* knockdown respectively decreased and increased anthocyanin biosynthesis and expression of the key TF MYB75 and of LBG-encoded target enzymes, such as DFR and LDOX (Baena-González et al., 2007, Nukarinen et al., 2016, Wang et al., 2021, Zirngibl et al., 2022). Repressed SnRK1 activity upon increased T6P levels is conversely characterized by induction of the flavonoid biosynthesis pathway (Zhang et al., 2009, Nunes et al., 2013). Mounting evidence suggests that SnRK1 regulation of flavonoid metabolism involves multiple levels of regulation. In addition to the transcriptional regulation of the LBGs, SnRK1 was shown to negatively regulate the upstream phenylpropanoid pathway via degradation of the PAL protein (Wang et al., 2021). Interestingly, SnRK1 was also shown to phosphorylate and inactivate HMG-CoA (3-hydroxy-3-methylglutaryl-CoA), the rate limiting enzyme in the cytosolic mevalonate pathway of terpenoid biosynthesis, another important source of specialized metabolites (Robertlee et al., 2017). Here we further explored the molecular mechanisms underlying SnRK1 regulation of anthocyanin production.

We first set up a simple seedling assay with mutant and transgenic plants to confirm the sucrose-specific induction and SnRK1 inhibition of anthocyanin biosynthesis through *MYB75* and subsequent target *LBG* expression (Figure 1, 2; Figure S1). To identify components in the signaling pathway controlling *MYB75* transcription in response to metabolic status, we previously evaluated the reported interactions of sucrose with hormone (and immune) signaling and identified the DELLA proteins as novel positive regulators in sucrose-induced anthocyanin biosynthesis. Sucrose specifically inhibits GA-mediated DELLA degradation (Li et al., 2014). In *tps1* mutants (deficient in T6P synthase) and SnRK1α1 OX plants (both showing reduced induction of anthocyanin biosynthesis), no consistent change in *GA3OX1* and *GA20OX1* GA/DELLA target gene expression could be observed (Li et al., 2014), suggesting that T6P–SnRK1 signaling might not be directly involved in sucrose-mediated DELLA stabilization as a mechanism to regulate *MYB75* expression. Several types of TFs have been reported to affect *MYB75* expression and anthocyanin biosynthesis, including B-Box binding (BBX) TFs, involved in many different aspects of photomorphogenesis through genetic and physical interaction with bZIP (HY5) and bHLH TFs, and TCP-type TFs (Shin et al., 2013; Li et al., 2015; Chang et al., 2008; Li and Zachgo, 2013; Viola et al., 2016). Nitrogen deficiency and sucrose induction of anthocyanin biosynthetic and regulatory genes is also mediated by histone acetylation (Liao et al., 2022). SnRK1 repression of *MYB75* expression may involve targeting any of those mechanisms. Finally, TAS4 (Trans-Acting SiRNA Gene 4)-siRNA81 is also targeting the mRNA of *MYB75* and *MYB90* post-transcriptionally in a sugar-responsive feedback loop (Luo et al., 2012).

The post-translational regulation of the MYB75 protein by SnRK1 described here adds another level to the regulation of anthocyanin biosynthesis. We investigated the mechanisms involved using cellular assays and a *DFR* promoter-LUC reporter construct. MYB75 functions in a heterotrimeric MBW complex (Zimmermann et al., 2004), but we already detected a 7-fold induction of *DFR* promoter activity upon transient overexpression of MYB75 in leaf mesophyll protoplasts, confirming its key role. However, expressing the MYB75-bHLH2 dimer produced a synergistic effect on promoter activity and analysis using *ttg1* KO protoplasts highlight the important function of the TTG1 WD40 domain protein for full transcriptional complex activity (Figure 3A-C). Most strikingly, we observed an SnRK1 phosphorylation dependent accelerated proteasomal degradation of MYB75 (Figure 5B, C). Such regulation was also reported for the Arabidopsis WRINKLED1 and Barley WRKY3 TFs (Zhai et al., 2017;Han et al., 2020). Anthocyanins accumulation is a photomorphogenic response and both MYB75 and MYB90 are degraded in dark conditions via ubiquitination by the COP1/SPA E3 ligase and 26S proteasome-dependent degradation (Li et al., 2012; Maier et al., 2013). In the presence of light, the COP1/SPA complex is repressed by photoreceptors (Podolec and Ulm, 2018). Recently, high light-induced anthocyanin biosynthesis was also reported to require inactivation of SnRK1 by increased sugar levels (Zirngibl et al., 2022), underscoring the physiological relevance of SnRK1 control.

We show that TTG1 and bHLH2 are also phosphorylated by SnRK1 (Figure 7B,D-F). While stability of the TTG1 protein did not alter upon SnRK1α1 co-expression, we did observe an increased cytosolic TTG1 localization (Figure 5E,F), consistent with complex dissociation. Indeed, co-IP assays indicate that SnRK1α1 decreases the interaction between the MBW subunits (MYB75-bHLH2, MYB75-TTG1 and bHLH2-TTG1) (Figure 5A). Whether the change in subcellular localization of TTG1 is linked with its phosphorylation state or a simple consequence of complex dissociation is unclear at the moment. Similarly, MYB75 degradation appears to be linked to complex dissociation as co-expression of complex members can increase MYB75 stability (Figure 5D). Finally, ChIP analysis indicates that SnRK1 also causes release of MYB75 from the *DFR* promoter chromatin (Figure 6D). Whether this is the cause or consequence of complex dissociation also remains to be explored. Our results also fit the hypothesis that TTG1 functions as a recruiter of bHLH to the promoter-bound R2R3-MYB TF, although another (complementary) function of TTG1 may be the shielding of the bHLH and R2R3-MYB factors from negative regulators (Zhang and Schrader, 2017).

We did not include the MYB123/TT2 (TRANSPARANT TESTA2) TF in our analyses as it is predominantly expressed and active as a key determinant for proanthocyanin accumulation in developing seeds (Nesi et al., 2001). However, the binding location of TT2 to the *DFR* promoter was studied earlier in *Physcomitrella patens* protoplasts, using overexpression of the *Arabidopsis* MYB123/TT2-bHLH42/TT8-TTG1 MBW complex and *Arabidopsis DFR* promoter fragments of various lengths with a GFP reporter (Xu et al., 2014). This study identified the 302 bp upstream of the translation start site as the minimal promoter containing the crucial regulatory elements to drive transcriptional activity. While we also observed a further decrease in *DFR* promoter activation by the MYB75-bHLH2-TTG1 MBW complex when truncating the 300 bp promoter sequence, our analyses pointed to a more important regulatory region between 350 and 300 bp (Figure 6A), containing a G-box-like bHLH recognition sequence (5’-CACGTC-3’), flanking a MYB-core element (5’-CACCAAAC-3’). Site-directed mutagenesis of these promoter elements significantly reduced the capacity of the MBW complex to induce *DFR* promoter activity (Figure 6C).

The activity of MBW complexes is known to be fine-tuned by competitive interaction with single-repeat R3 MYBs (Zimmermann et al., 2004; Matsui et al., 2008; Wang et al., 2008; Zhu et al., 2009). CAPRICE (CPC), for example, affects root hair differentiation, trichome formation, stomatal development, and possibly also anthocyanin biosynthesis (Wada et al., 1997; Zhu et al., 2009). *Arabidopsis* plants ectopically overexpressing MYBL2 are characterized by repressed leaf trichome development and decreased anthocyanin accumulation (Sawa, 2002; Matsui et al., 2008). However, while *mybl2* KO mutants show an increase in leaf anthocyanin levels, this mutant still has a normal trichome phenotype (Dubos et al., 2008; Matsui et al., 2008). We confirmed the repressive effect of MYBL2 on MBW complex activity in transient overexpression assays with leaf mesophyll protoplasts and identified SnRK1 as a positive regulator of *MYBL2* expression (Figure 8A-C). This is consistent with an earlier observation of decreased *MYBL2* mRNA transcript levels in high light (increasing sugar content) conditions (Dubos et al., 2008; Zirngibl et al., 2022).

Anthocyanin biosynthesis and accumulation have been used as obvious and convenient phenotypes in serendipitous discoveries as well as in the development, validation, and optimization of genetic and molecular tools, such as RNAi, CRISPR-Cas mediated gene editing, activation tagging, or EAR motif-mediated transcriptional repression (Napoli et al., 1990; van der Krol, 1990; Borevitz et al., 2000; Numata et al., 2014; Kiselev et al., 2021; Khusnutdinov et al., 2021). Here, we describe that anthocyanin biosynthesis also reflects and acts as a physiologically relevant readout of plant metabolic status and SnRK1 activity. However, there is also natural variation in sucrose-induced anthocyanin accumulation and the Cvi ecotype is almost non-responsive (Teng et al., 2005), due to a mutation in the MYB75 protein (Figure S3A). This ecotype originates from the Cape Verde Islands close to the equator and is well adapted to drought, high temperatures and high irradiance levels. Given its high photooxidative stress tolerance, the lack of anthocyanin biosynthesis is surprising. Apparently, these plants have evolved alternative protective systems, including a unique chloroplastic copper/zinc superoxide dismutase (Abarca et al., 2001).

Combining our results, we propose a model in which SnRK1 inhibits the MBW complex controlling anthocyanin biosynthesis at both the transcriptional and post-translational level (Figure 8D). Transcriptional regulation involves repression of *MYB75* expression and induction of *MYBL2* expression, either directly or indirectly. Post-translational regulation by SnRK1 involves MBW complex phosphorylation and dissociation and (subsequent) MYB75 protein degradation and TTG1 translocation. Detailed mechanistic insight into how exactly anthocyanin biosynthesis is affected by plant metabolic status might not only enable the identification of better molecular markers but also the uncoupling of anthocyanin biosynthesis from metabolic stress regulation through breeding and engineering for increased plant protection, food quality, and human health. However, the extensive regulation revealed in this study indicates that repression of anthocyanin biosynthesis, and possibly specialized metabolism more generally, is an important strategy to save energy and redirect C flow to more essential processes for survival in metabolic stress conditions.

## Acknowledgments

We are grateful to María Rosa Ponce Molet for *MYB75 RNAi* mutant seeds, Chris Lamb for *pap1-D* seeds, Maarten Koornneef for *ttg1-1* (*W89*) seeds, Markus Schmid for the *tps1-2 GVG:TPS1* suppressor mutants, and the Nottingham Arabidopsis Stock Centre (NASC) for *dfr1, ttg1-21* (GK-580A05) and *ttg1-22* (GK-286A06) mutant seeds. Anja Vandeperre and Hilde Verlinden provided excellent technical assistance and plant care.

## Materials and Methods

### Plant Materials and Growth Conditions

*Arabidopsis thaliana* mutant lines were in the Col-0 or Ler background. The Ler *SnRK1α1 OX, SnRK1α1 RNAi* (Baena-González et al., 2007), *dfr1* (*tt3-1*; NASC NW84) and Col-0 *pap1-D* (Borevitz et al., 2000), *myb75 RNAi* (Jover-Gill et al., 2014), *ttg1-21* (GK-580A05; NASC N2105595), *ttg1-22* (GK-286A06; NASC N330696), *tps1* (Van Dijken et al., 2004), *tps1* suppressor (Zacharaki et al., 2022) and *NLS-SnRK1α1 and βMYR-SnRK1α1* (Ramon et al., 2019) mutants and transgenic lines have been described previously.

Seeds were vapor-sterilized and stratified for 3 days at 4°C. For each biological replicate, 15-20 seeds were germinated and grown in 6-well plates in liquid half-strength Murashige and Skoog (MS) medium supplemented with 0.5 % (w/v) glucose. Seeds were incubated under continuous white fluorescent light (65uE) at 21°C.

For anthocyanin measurements, liquid medium was exchanged after 4 days by liquid half-strength MS medium supplemented with either 100 mM mannitol (osmotic control), 100 mM glucose, or 100 mM sucrose. Seedlings were incubated another 3 days before anthocyanin levels were quantified.

For gene expression analysis, liquid medium was replaced after 7 days by liquid half-strength MS medium supplemented with either 100 mM mannitol, 100 mM glucose, or 100 mM sucrose. Seedlings were snap frozen in liquid nitrogen 6 hours after the medium was replaced to study short term transcriptional sugar effects via qRT-PCR.

For the starvation assay, liquid medium was replaced after 7 days by liquid half-strength MS medium without any carbon source. Seedlings were snap frozen in liquid nitrogen after 6 hours sugar starvation to study short term transcriptional responses via qRT-PCR.

### Anthocyanin quantification

Anthocyanins were extracted and quantified as described by Rabino and Mancinelli (1986). 7-day-old seedlings were harvested, weighed, and incubated in 1 mL extraction buffer (acidic methanol, 1 % HCl w/v) for 24 hours at 4°C in the dark. Absorbance of the supernatants was measured at 530 nm and 657 nm (NanoPhotometer™, Implen). Relative anthocyanin levels were quantified as (A_350nm_-1/4A_657nm_) per gram fresh weight.

### Plasmid construction

Full length *Arabidopsis MYB75* (*At1g56650*), *bHLH2/EGL3* (*At1g63650*), *TTG1* (*At5g24520*), *MYBL2* (*AT1G71030*) and *SnRK1α1/KIN10* (*At3g01090*) coding sequences without the stop codon were PCR-amplified from *Arabidopsis* Col cDNA and inserted in the pUC18-based HBT95 overexpression vector containing the 35SC4PPDK promoter (35S enhancer and maize C4PPDK basal promoter) and the nopaline synthase (NOS) terminator, in-frame with a double HA-, FLAG- or with an eGFP-tag (Sheen, 1996).

The *DFR* (*At5g42800*) promoter sequences were PCR-amplified from genomic DNA and ligated in a pUC18-based vector in front of the firefly luciferase (LUC) reporter gene (Yoo et al., 2007). Point mutations were made via site-directed PCR mutagenesis using complementary mutant primer pairs extending 15 nucleotides on either side of the modification. Methylated template DNA was digested with *DpnI*.

### Transient Expression in Leaf Mesophyll Protoplasts

Isolation and PEG/Ca^2+^-mediated transfection of *Arabidopsis* leaf mesophyll protoplast was performed as described in Yoo et al. (2007). Transfected cell volumes (with 20,000 cells/mL) varied depending on the experiment (50 μL for LUC assays, 100 μL for immunoblot assays, 1 mL for co-IP experiments, 1.5 mL for qRT-PCR expression analyses and 3 mL for Chip assays). CsCl gradient purified plasmid DNA was used to transfect the cells. Transfected cells were exposed to dim light and harvested after 6 hours of incubation for LUC and qRT-PCR assays. For protein stability assays, 10 μM cycloheximide was added to the protoplasts 4 hours after transfection. A sample was harvested every hour for immunoblot analysis. Protoplasts were harvested by centrifugation at 1100 rpm (200 g) using a swinging bucket rotor (5804 Eppendorf) and the pellets were stored at −80°C.

### LUC and GUS Assays

Transfected protoplasts were lysed with 50 μL lysis buffer (25 mM Trip-Phosphate at pH 7.8, 2 mM DTT, 2 mM 1.2-diaminocyclohexane-N, N,N’,N’-tetra-acetic acid, 10% [v/v] glycerol, and 1% [v/v] Triton X-100). 100 μL LUC assay reagent (E1500 Kit; Promega) was added to 20 μL cell lysate into a luminometer tube. A Lumat LB9507 tube luminometer (Berthold Technologies) was used to detect luminescence.

Correction for pipetting errors and transfection efficiency was done using a co-transfected UBQ-GUS construct. 5 μL of cell lysate was added to 45 μL of 10 mM 4-methylumbelliferyl-β-D-glucuronide solution (MUG, M-9130; Sigma-Aldrich). The reaction was stopped after 1 hour incubation at 37°C by adding 220 μL of 0.2 M Na_2_CO_3_. Fluorescence was measured with the GloMax^-^Multi^+^ Detection System (Promega).

### qRT-PCR

TRIsure™ Reagent (Bioline) was used to extract RNA from seedlings, rosette leaves and transfected protoplasts, following manufacturer’s instructions. Isolated RNA was converted into cDNA using the SensiFAST™ cDNA synthesis kit (Bioline). Quantification of the relative amount of specific mRNAs was done using quantitative real time PCR in a 96-well plate with the StepOne™ Real PCR system. PowerUp™ SYBR™ Green Master mix was used to perform qRT-PCR (Thermo Fisher Scientific). 10 ng cDNA was mixed in a total volume of 10 μL with 5 μL PowerUp SYBR Green master mix, 0.2 μL of each primer and 2.6 μL H_2_O. Thermal cycling conditions used: 2 minutes at 50°C, 2 minutes at 95°C and 40 cycles of 3 seconds at 95°C and 30 seconds at 60°C. Marker gene expression was normalized to *UBQ10* or *eIF4A* gene expression, chosen based on their stable expression in different tissues and metabolic stress conditions (Czechowski et al., 2005).

### Immunoblot Analyses

Transiently overexpressed proteins were detected using immunoblot analysis. Before loading on a polyacrylamide gel, loading buffer (120 mM Tris-HCl at pH 6.8, 5.4 M urea, 20% [v/v] glycerol, 4% [w/v] SDS, 5% [v/v] β-mercaptoethanol, and 0.5% [v/v] bromophenol blue) was added to the samples, followed by heating at 95°C for 5 minutes. Proteins were separated on a 1.5 mm 10 % acrylamide SDS-PAGE gel in Tris-Gly running buffer (25 mM Tris, 192 mM Gly, and 0.1 % [w/v] SDS at pH 8.5). Separation was obtained through stacking for 15 minutes at 60 V and 15 minutes at 110 V followed by running for 1 hour at 160 V. Separated proteins were transferred from the gel to a polyvinylidene fluoride membrane (PVDF, Immobilon-P Millipore) using a wet blot system (Mini-PROTEAN Tetra Cell, Bio-Rad) in Tris-Gly buffer with 10 % (v/v) methanol for 2 h at 300 mA. After blotting, the membrane was blocked for 1 h with 5 % skimmed milk, followed by a 2 h incubation at RT (or overnight at 4°C) with Horse Radish Peroxidase (HRP)-conjugated antibody (HRP-conjugated anti-HA antibody, 1/1000 [50 mg/mL], cat. no. 12013819001, Roche; HRP-conjugated anti-FLAG antibody, 1/1,000 [1 mg/mL], cat. no. A8592, Sigma-Aldrich). The membrane was washed three times (TBST; 500mM NaCl, 13.5mM Tris-HCl pH 7.5 and 0.05% Tween®20). Antibody-bound proteins were detected by incubating the membrane in the Supersignal® West Pico Plus Chemiluminescent substrate (cat. no. 34,577; Thermo Fisher Scientific) for 2 minutes and visualized by exposure to light sensitive film (CL-XPosureTM Film 5×7 inches, ThermoScientific). Ribulose bisphosphate carboxylase small chain (RBCS) staining of the blot with Coomassie Brilliant Blue R-250 was used as a protein loading control.

### Phosphorylation Mobility Shift Assay

A phosphate-binding compound (Phos-tag, Wako Chemicals; cat. no. AAL-107) and MnCl_2_(H_2_O)4 was added to the 1.5 mm 10 % (w/v) poly-acrylamide gel to allow visualization of a mobility shift of phosphorylated proteins. Protein samples were prepared by adding loading buffer and heating for 5 min at 95°C. Separation was obtained running at 30 mA. Before blotting, the gel was soaked two times in transfer buffer containing 10% (v/v) methanol and 10 mM EDTA for 20 minutes with gentle agitation, followed by 20 minutes in transfer buffer without EDTA. The next steps, transferring the proteins to a PVDF membrane and visualization, were as described for “Immunoblot Analyses”.

### Subcellular localization

Subcellular localization of transiently overexpressed proteins was observed using confocal laser scanning microscopy (FV1000; Olympus). Protoplasts were transfected with eGFP construct DNA and incubated for 6 to 16 hours prior to visualization.

### Co-immunoprecipitation

Leaf mesophyll protoplasts transiently co-expressing HA- and FLAG-tagged recombinant proteins were lysed with 200 μL co-IP buffer containing protease inhibitor [50 mM Tris-HCL pH7.5, 150mM NaCl, 5mM EDTA, 1% Triton X-100, 0.5 mM DTT, 1x Protease inhibitor cocktail]. A 20 μL aliquot (input control) was immediately frozen at −80°C. 20 μL of anti-FLAG antibody-couple agarose beads (Sigma-Aldrich) were added to the remaining cell lysate for overnight rotating incubation at 4°C. After incubation, beads were washed at least three times with 500 μL co-IP buffer without protease inhibitor. Eluted samples and input samples were subjected to immunoblot analysis using conjugated anti-HA and anti-FLAG antibodies.

### Chromatin Immunoprecipitation (ChIP)

The ChIP assay was performed as previously described (Nelson et al., 2006). Wildtype leaf mesophyll *Arabidopsis* protoplasts were transfected with the HA-tagged constructs of interest (150 μL DNA per 3 mL protoplasts) and incubated for 6 hours. Samples were crosslinked with 1 % formaldehyde, quenched with glycine, and washed with cold TBS buffer. Nuclei were isolated from the crosslinked samples and chromatin was sheared via sonication using a Bioruptor sonicator (25 minutes, high power, 30 sec on/ 30 sec off). Sonicated lysates were cleared via centrifugation at maximum speed for 10 min at 4 °C. Half of the supernatants was used as input DNA sample in which the DNA was precipitated with 100 % ethanol, the other half for immunoprecipitation (IP). For IP, anti-HA antibody was added to the samples and rotated overnight at 4°C. The protein-chromatin complex was captured using 40 μL slurry protein G agarose beads and rotation for 1 hour. The beads were washed, and chromatin was isolated from both input and IP samples using 10 % (w/v) Chelex 100. Proteases were inactivated through boiling the samples with 20 μg proteinase K for 1 hour at 50°C. The samples were centrifuged and the chromatin containing supernatants was transferred into a new tube. ChIP products of both input and IP samples were analyzed via qRT-PCR with *UBQ10* as background control. After calculating the signal ratio, relative enrichment of target regions were normalized against *UBQ10*. The ChIP experiments were performed with three biological replicates, from which the means and standard deviations were calculated.

### In vitro kinase assay

Gateway ORF entry clones of SnRK1α1, SnRK1α1K48M, MYB75, TTG1, TTG1S94A and bHLH2 were cloned into pDest-HisMBP through standard LR gateway reaction. The resulting His-MBP expression vectors were transformed into *E. coli* BL21 for production of recombinant proteins as previously described (Van Leene et al., 2019). Radioactive kinase reactions were performed in kinase assay buffer (50 mM Tris-HCl (pH 8.0), 1 mM EGTA, 1 mM DTT, 5 mM MgCl_2_, 10 μM cold ATP, 5 μCi γ-^32^P ATP, 1x PhosSTOP) for 1 h at 30°C, combining 2 μL kinase with 4-15 μL substrate. Amicon-purified MBP elution buffer was added to correct for varying amounts of recombinant proteins in each reaction. Reactions were stopped by addition of SDS sample buffer and incubation for 10 min at 95°C. For detection of radiolabeled phosphoproteins, proteins were separated by SDS-PAGE on TGX 4-15 % gradient gels (Biorad) and stained with Coomassie brilliant blue R-250. Gels were dried and radioactivity was detected by autoradiography on a photographic film.

For mass spectrometry-based identification of phosphopeptides, kinase assays were performed as described above, using 10 μM cold ATP instead of γ-^32^P ATP and reactions were incubated overnight at 30°C. Reactions were stopped by addition of NuPAGE sample buffer (ThermoFisher Scientific) and incubation at 70°C for 10 min. Proteins were separated for 7 min at 200 V on a 4-12 % NuPAGE gradient gel, stained with Coomassie G-250 and in-gel trypsin digested (Van Leene et al., 2015). Peptides were re-dissolved in 15 μL loading solvent A (0.1 % TFA in water/ACN (98:2, v/v)) of which 5 μL was injected for LC-MS/MS analysis on an an Ultimate 3000 RSLC nano LC (Thermo Fisher Scientific, Bremen, Germany) in-line connected to a Q Exactive mass spectrometer (Thermo Fisher Scientific). The peptides were first loaded on a μPAC™ Trapping column with C18-endcapped functionality (Pharmafluidics, Belgium) and after flushing from the trapping column the peptides were separated on a 50 cm μPAC™ column with C18-endcapped functionality (Pharmafluidics, Belgium) kept at a constant temperature of 35°C. Peptides were eluted by a linear gradient from 98 % solvent A’ (0.1 % formic acid in water) to 55 % solvent B’ (0.1 % formic acid in water/acetonitrile, 20/80 (v/v)) in 30 min at a flow rate of 300 nL/min, followed by a 5 min wash reaching 99% solvent B’. The mass spectrometer was operated in data-dependent, positive ionization mode, automatically switching between MS and MS/MS acquisition for the 5 most abundant peaks in a given MS spectrum. The source voltage was 3.0 kV, and the capillary temperature was 275°C. One MS1 scan (m/z 400-2,000, AGC target 3 × 106 ions, maximum ion injection time 80 ms), acquired at a resolution of 70,000 (at 200 m/z), was followed by up to 5 tandem MS scans (resolution 17,500 at 200 m/z) of the most intense ions fulfilling predefined selection criteria (AGC target 5 × 104 ions, maximum ion injection time 80 ms, isolation window 2 m/z, fixed first mass 140 m/z, spectrum data type: centroid, intensity threshold 1.3xE4, exclusion of unassigned, 1, 5-8, >8 positively charged precursors, peptide match preferred, exclude isotopes on, dynamic exclusion time 12 s). The raw MS files were processed with the MaxQuant software (version 1.6.4.0) (Cox and Mann, 2008), and searched with the built-in Andromeda search engine against the Araport11plus database. This database consists of the Araport11 database with non-plant common Repository of Adventitious Proteins (cRAP) sequences e.g. tags, keratins, trypsin etc. added. MaxQuant search parameters and MS results can be found in Supplemental Table S3.

