## Supplementary Data for "SnRK1 inhibits anthocyanin biosynthesis through both transcriptional regulation and direct phosphorylation and dissociation of the MYB/bHLH/TTG1 MBW complex"

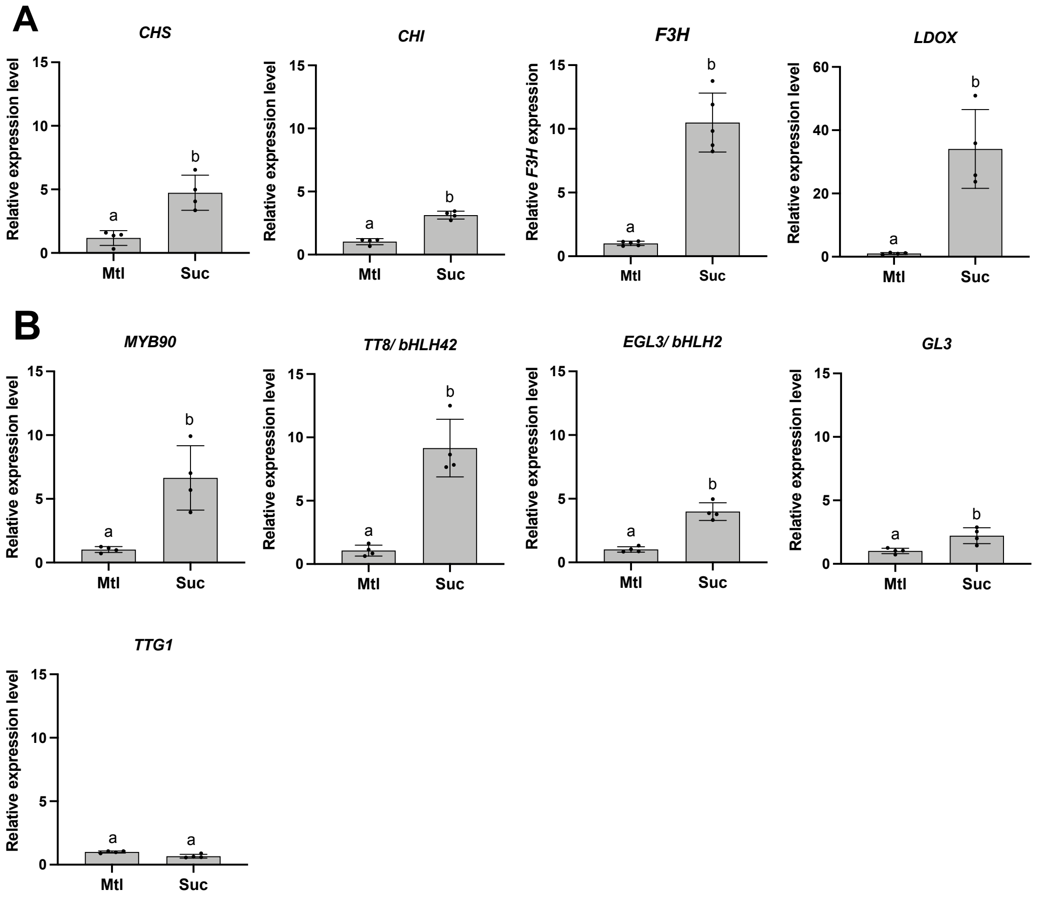


***Supplementary Figure 1. Sucrose induction of anthocyanin biosynthesis-related enzyme and transcription factor gene expression.***

*qRT-PCR analysis of gene expression levels of (A) three upstream flavonoid biosynthesis enzymes (CHS, CHI, F3H) and one more downstream biosynthetic enzyme (LDOX), and (B) regulatory proteins (MYB90, TT8/bHLH42, EGL3/bHLH, GL3, TTG1) in 7-day-old Col-0 wildtype seedlings grown in ½ MS medium supplemented with 100 mM mannitol or 100 mM sucrose. Values are averages with SD, n = 4 biological repeats. Unpaired t-test analysis was performed in GraphPad Prism v9, letters represent statistically significant differences, p<0.012.*

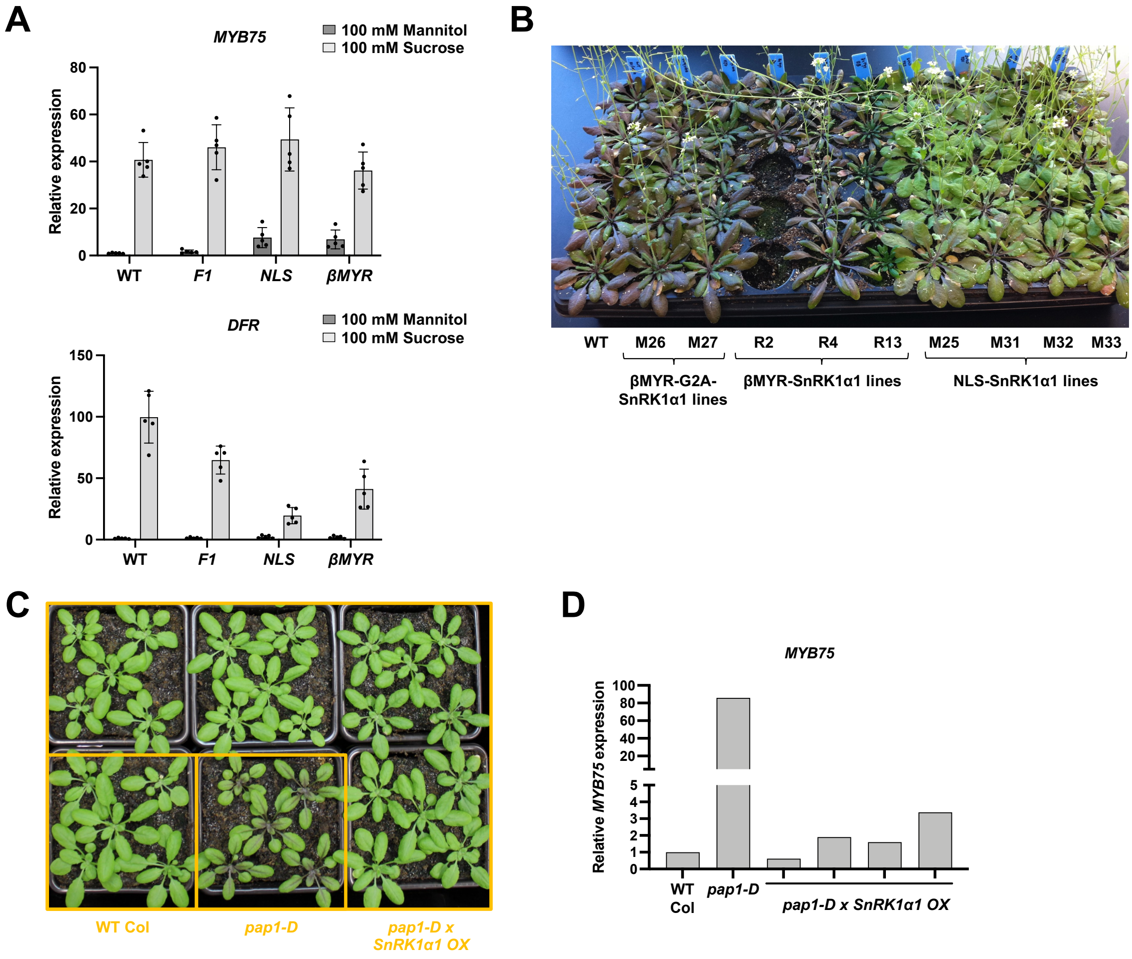


***Supplementary Figure 2. Overexpression of SnRK1α1 in the pap1-D mutant.***

*(A) qRT-PCR analysis of MYB75 and DFR gene expression levels 7-day-old Col-0 wildtype and SnRK1α1/ SnRK1α1 SnRK1α2/ SnRK1α2 double mutant seedlings complemented with wildtype SnRK1α1 (F1), NLS-SnRK1α1 (NLS), and βMYR-SnRK1α1 (βMYR) seedlings grown in ½ MS medium supplemented with 100 mM mannitol or 100 mM sucrose. Values are averages with SD, n = 5 biological repeats.*

*(B) Phenotype of Col-0 wildtype plants, 2 independent βMYR-G2A-SnRK1α1 lines (M26, M27), 3 independent βMYR-SnRK1α1 lines (R2, R4, R13) and 4 independent NLS-SnRK1α1 lines (M25, M31, M32, M33).*

*(C) Phenotype of 4-weeks-old soil-grown Col-0 wildtype, pap1-D and pap1-D x SnRK1α1 OX plants.*

*(D) qRT-PCR analysis of MYB75 gene expression levels in 4-weeks-old soil-grown Col-0, pap1-D and pap1-D x SnRK1α1 OX plants.*

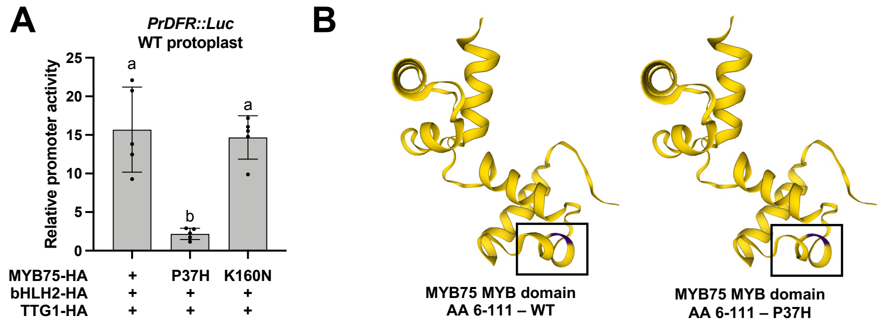


***Supplementary Figure 3. Activity of MYB75 in DFR transcription upon introduction of Cvi ecotype-specific mutations.***

*(A) DFR promoter activity in Arabidopsis leaf mesophyll protoplasts upon transient co-expression of bHLH2, TTG1 and wildtype or mutated (P37H, K160N) MYB75. Relative and normalized promoter activity values are averages with SD, n = 5 biological repeats (independent protoplast transfections). One-way ANOVA statistical analysis was performed in GraphPad Prism v9, letters represent statistically significant differences, p<0.0005.*

*(B) Protein 3D structure prediction of the MYB domain of WT Col MYB75 (left) and mutated MYB75 P37H (right). Modelling was done in Swiss-Model based on the crystal structure of the well-characterized R2R3 MYB protein MYB domain WER. The residues P37 and H37 are indicated in purple.*

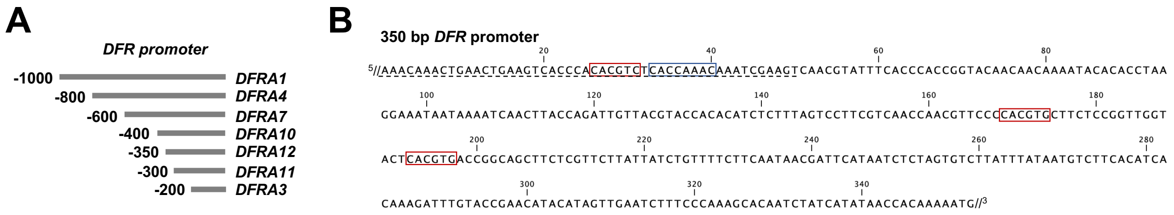


***Supplementary Figure 4. DFR promoter dissection.***

*(A) Schematic representation of the truncated DFR promoter fragments: 1000 bp (DFRA1), 800bp (DFRA4), 600bp (DFRA7), 400bp (DFRA10), 350bp (DFRA12), 300 bp (DFRA11), and 200bp (DFRA3).*

*(B) DFR promoter sequence 350bp upstream of the ATG start codon. The G-box-like and two perfect CACGTG G-boxes are highlighted with red frames, the putative MYB core element is highlighted with a blue frame. The 50 bp stretch is underlined.*

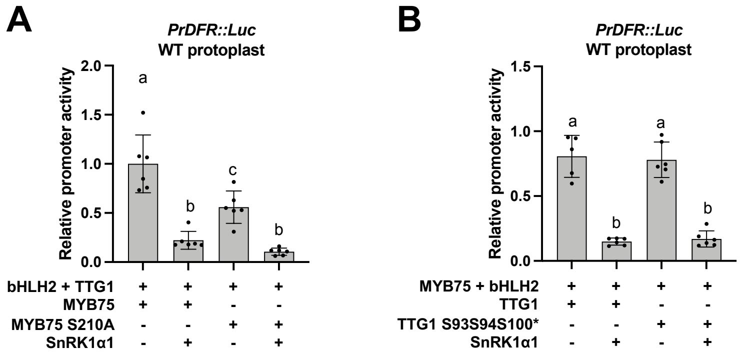


***Supplementary Figure 5. Effect of MYB75 and TTG1 phosphorylation on MBW complex activity.***

*(A) DFR promoter activity in leaf mesophyll protoplasts transiently expressing the MBW complex with wildtype or mutated TTG1 or MYB75 proteins without or with co-expression of SnRK1α1. Values are averages with SD, n = 5-6 biological repeats (independent protoplast transfections). One-way ANOVA statistical analysis was performed in GraphPad Prism v9, letters represent statistically significant differences, p<0.05.*

*Supplementary Table 1: Cloning and mutagenesis primers.*

| ***Gene/promoter*** | ***Use*** | ***Primer sequence*** |
|---|---|---|
| *MYB75* | *Cloning CDS* | *CG-GGATCC-ATGGAGGGTTCGTCCAAAG*<br>*A-AGGCCT-ATCAAATTTCACAGTCTCTCC* |
| *bHLH2* | *Cloning CDS* | *CG-GGATCC-ATGGCAACCGGAGAAAACAG*<br>*A-AGGCCT-ACATATCCATGCAACCCTTTG* |
| *TTG1* | *Cloning CDS* | *CG-GGATCC-ATGGATAATTCAGCTCCAGAT*<br>*A-AGGCCT-AACTCTAAGGAGCTGCATTTT* |
| *MYBL2* | *Cloning CDS* | *GA-AGATCT-ATGAACAAAACCCGCCTTCGT*<br>*A-AGGCCT-TCGGAATAGAAGAAGCGTTTCTT* |
| *DFR* | *Cloning promoter* | *CG-GGATCC-CTTTGTCTCTCTGTTTGGAG*<br>*CATG-CCATGG-TTTTGTGGTTATATGATAGATTGT* |
| *Prom DFR-A4* | *Cloning promoter* | *CG-GGATCC-ATTTCTTACTTTATGAGATTAAGT* |
| *Prom DFR-A7* | *Cloning promoter* | *CG-GGATCC-ATGTTTTATAATTTTACAATTTTGT* |
| *Prom DFR-A10* | *Cloning promoter* | *CG-GGATCC-GTTTAGACAAATTTTGATTTATTTCG* |
| *Prom DFR-A12* | *Cloning promoter* | *CG-GGATCC-AAACAAACTGAACTGAAGTCACC* |
| *Prom DFR-A11* | *Cloning promoter* | *CG-GGATCC-CAACGTATTTCACCCACCGG* |
| *Prom DFR-A3* | *Cloning promoter* | *CG-GGATCC-TCGTCAACCAACGTTCCCC* |
| *SnRK1α1* | *Cloning CDS* | *CG-GGATCC-ATGGATGGATCAGGCACAGGC*<br>*A-AGGCCT-GAGGACTCGGAGCTGAGCAA* |
| *SnRK1α1 K48M* | *SDM* | *CATAAGGTTGCTATCATGATCCTCAATCGTCGC*<br>*GCGACGATTGAGGATCATGATAGCAACCTTATG* |
| *MYB75 T130A* | *SDM* | *CGCCCATTCCTACAGCACCGGCACTAAAAAAC*<br>*GTTTTTTAGTGCCGGTGCTGTAGGAATGGGCG* |
| *MYB75 T126AT130AT131A* | *SDM* | *GAAAAAGAGAGACATTGCGCCCATTCCTGCAGCACCGGCACTAAAAAAC*<br>*GTTTTTTAGTGCCGGTGCTGCAGGAATGGGCGCAATGTCTCTCTTTTTC* |
| *MYB75 S210A* | *SDM* | *AAATTCCTAGAGGAAGCCCAAGAGGTAGATATT*<br>*AATATCTACCTCTTGGGCTTCCTCTAGGAATTT* |

| | | |
|---|---|---|
| *TTG1 T14A* | *SDM* | *GTTATCCAGATCGGAAGCCGCCGTCACATACGAC*<br>*GTCGTATGTGACGGCGGCTTCCGATCTGGATAAC* |
| *TTG1-S93A* | *SDM* | *TCTCTCCGTCGTCCTGCCTCCGGAGATCTCCTC*<br>*GAGGAGATCTCCGGAGGCAGGACGACGGAGAGA* |
| *TTG1 S94A* | *SDM* | *CTCCGTCGTCCTTCCGCCGGAGATCTCCTCGCTTC*<br>*GAAGCGAGGAGATCTCCGGCGGAAGGACGACGGAG* |
| *TTG1 S100A* | *SDM* | *GGAGATCTCCTCGCTGCCTCCGGCGATTTCCTC*<br>*GAGGAAATCGCCGGAGGCAGCGAGGAGATCTCC* |
| *TTG1 S165A* | *SDM* | *GGGATATTGAGAAGGCTGTTGTTGAGACTCAG*<br>*CTGAGTCTCAACAACAGCCTTCTCAATATCCC* |
| *TTG1 S93AS94A* | *SDM* | *TCTCTCCGTCGTCCTGCCGCCGGAGATCTCCTCGCT*<br>*AGCGAGGAGATCTCCGGCGGCAGGACGACGGAGAGA* |

*Supplementary Table 2: qRT-PCR primers.*

| *Gene* | *Forward* | *Reverse* |
|---|---|---|
| *DFR* | *CTTTGTTCGTGCCACCGTTCG* | *TCCTTCCTCAGATAAATCAGCCTTCC* |
| *MYB75* | *GGAATAGGTGGTCTTTAATTGCTG* | *AGGAATGGGCGTAATGTCTC* |
| *UF3GT* | *TGTCAGATCGTTTTGGTTCC* | *GATTCTTCCTCACTTTCTCAC* |
| *LDOX/ANS* | *GTTTGCAGCTTTTCTACGAGG* | *TGAGCAAAAGTCCGTGGAGG* |
| *CHS* | *GGCAAAGAAGCGGCAGTGAAG* | *CGGAAGGACGGAGACCAAGAAG* |
| *CHI* | *GGAGGCGGTTCTGGAATCTATC* | *TCGTCCTTGTTCTTCATCATTAGC* |
| *FLS* | *CCACCGTCATGCGTCAATTACAG* | *TCTCCGCCGAGACCTTCTTTCAA* |
| *TT8* | *GGCGGTTCAATCTGTGGAC* | *CTGTTGGCTCCTCTCTAACG* |
| *bHLH2/EGL3* | *GGATTCAACGGTTAGGTCAG* | *ATCATCTTCTGCGATTTCTCTC* |
| *MYB90* | *AAGCTGATGCGATTGTTCCT* | *CCAAAGTTGCTCAACGTCAA* |
| *TT7 F3’H* | *TAGCCGACCACCAAACTC* | *AGCGTTCCAACCTCTTCC* |
| *TT6 F3H* | *TCGTCTCTAGTCACCTCCAG* | *TCACTTTCACCCAACCTTCC* |
| *UBQ10* | *AACTTTGGTGGTTTGTGTTTTGG* | *TCGACTTGTCATTAGAAAGAAAGAGATAA* |
| *eIF4A* | *GAAATCGCTTCCTTCGTTTG* | *ATAACTGCCAGCACCACCTC* |
| *MYBL2* | *TGCACTTCTTGGCAATAGATGGTC* | *TTGTCGGTTCGTCCTGGCAATC* |
| *DFR-CH1* | *AAAAACAAACTGAACTGAAGTCA* | *GTTGATTTTATTATTTCCTTAGGT* |
| *DFR-CH2* | *CAAATCGAAGTCAACGTATTTC* | *TGGTTGACGAAGGACTAAAGA* |
| *DFR-CH3* | *ACCTAAGGAAATAATAAAATCAA* | *AAGAAAACAGATAATAAGAACGA* |
| *DFR-CH4* | *ACGTTCCCCACGTGCTTCT* | *AGACATTATAAATAAGACACTAGA* |
| *DFR-CH5* | *CCAAAGCACAATCTATCATATA* | *CACGAACAAAGTAACCACGT* |
| *DFR-CH6* | *TTCGTAAAAGTGGGTGGGGA* | *CCGGTGGGTGAAATACGTTG* |
| *DFR-CH8* | *TTTAGATTGCAATAAATCTAGAAGTCA* | *TCTTGGAAAAGCTTAAATGAAAAA* |
| *DFR-CH9* | *TTTCTTTGCGAAAACTGGAAA* | *CAATTGTGTCCCATGTCAGC* |
| *DFR-CH10* | *AAGCTGACCTCTTCTCTGACG* | *ATTTCCAGTTTTCGCAAAGAAA* |
| *DFR-CH11* | *TCTCTGTTTGGAGGAGAGTCAA* | *CTCGCGTCAGAGAAGAGGT* |

*Supplementary Table 3: MaxQuant phosphosearch parameters.*

| ***Parameter*** | ***Value*** |
|---|---|
| *Version* | *1.6.4.0* |
| *Include contaminants* | *FALSE* |
| *PSM FDR* | *0,01* |
| *PSM FDR Crosslink* | *0,01* |
| *Protein FDR* | *0,01* |
| *Site FDR* | *0,01* |
| *Use Normalized Ratios For Occupancy* | *TRUE* |
| *Min. peptide Length* | *7* |
| *Min. score for unmodified peptides* | *0* |
| *Min. score for modified peptides* | *40* |
| *Min. delta score for unmodified peptides* | *0* |
| *Min. delta score for modified peptides* | *6* |
| *Min. unique peptides* | *1* |
| *Min. razor peptides* | *1* |
| *Min. peptides* | *1* |
| *Use only unmodified peptides and* | *TRUE* |
| *Modifications included in protein quantification* | *Oxidation (M);Acetyl (Protein N-term);Phospho (STY)* |
| *Peptides used for protein quantification* | *Unique* |
| *Discard unmodified counterpart peptides* | *TRUE* |
| *Label min. ratio count* | *1* |
| *Use delta score* | *FALSE* |
| *iBAQ* | *TRUE* |
| *iBAQ log fit* | *TRUE* |
| *Match between runs* | *TRUE* |
| *Matching time window [min]* | *0,7* |
| *Match ion mobility window [indices]* | *0,05* |
| *Alignment time window [min]* | *20* |
| *Alignment ion mobility window [indices]* | *1* |
| *Find dependent peptides* | *FALSE* |

| | |
|---|---|
| *Fasta file* | *Araport11plus_DE20191003.fasta* |
| *Decoy mode* | *revert* |
| *Include contaminants* | *FALSE* |
| *Advanced ratios* | *TRUE* |
| *Second peptides* | *TRUE* |
| *Stabilize large LFQ ratios* | *TRUE* |
| *Separate LFQ in parameter groups* | *FALSE* |
| *Require MS/MS for LFQ comparisons* | *TRUE* |
| *Calculate peak properties* | *FALSE* |
| *Main search max. combinations* | *200* |
| *Advanced site intensities* | *TRUE* |
| *Max. peptide mass [Da]* | *4600* |
| *Min. peptide length for unspecific search* | *8* |
| *Max. peptide length for unspecific search* | *25* |
| *Razor protein FDR* | *TRUE* |
| *Disable MD5* | *FALSE* |
| *Max mods in site table* | *3* |
| *Match unidentified features* | *FALSE* |
| *Evaluate variant peptides separately* | *TRUE* |
| *Variation mode* | *None* |
| *MS/MS tol. (FTMS)* | *20 ppm* |
| *Top MS/MS peaks per Da interval. (FTMS)* | *12* |
| *Da interval. (FTMS)* | *100* |
| *MS/MS deisotoping (FTMS)* | *TRUE* |
| *MS/MS deisotoping tolerance (FTMS)* | *7* |
| *MS/MS deisotoping tolerance unit (FTMS)* | *ppm* |
| *MS/MS higher charges (FTMS)* | *TRUE* |
| *MS/MS water loss (FTMS)* | *TRUE* |
| *MS/MS ammonia loss (FTMS)* | *TRUE* |
| *MS/MS dependent losses (FTMS)* | *TRUE* |
| *MS/MS recalibration (FTMS)* | *FALSE* |

### *Supplementary Table 4: SnRK1α1 Phospho(STY)Sites in TTG1.*

| **Positions within protein** | *S93* | *S94* | *S100* | *S12* | *S165* | *T14* | *T169* |
|---|---|---|---|---|---|---|---|
| **Localization prob** | *0.986154* | *0.5826* | *0.970153* | *0.5* | *0.854783* | *0.906818* | *0.624798* |
| **Sequence window** | *PPTKLMFSPPSLRRPSSGDLLASSGDFLRLW* | *PTKLMFSPPSLRRPSSGDLLASSGDFLRLWE* | *SPPSLRRPSSGDLLASSGDFLRLWEINEDSS* | *____MDNSAPDSLSRSETAVTYDSPYPLYAM* | *TCSIDTTCTIWDIEKSVVETQLIAHDKEVHD* | *__MDNSAPDSLSRSETAVTYDSPYPLYAMAF* | *DTTCTIWDIEKSVVETQLIAHDKEVHDIAWG* |
| **Phospho (STY) Probabilities** | *RPS(0.986)S(0.014)GDLLASSGDFLR* | *RPS(0.414)S(0.583)GDLLAS(0.003)SGDFLR* | *RPS(0.003)S(0.003)GDLLAS(0.97)S(0.023)GDFLR* | *S(0.5)ET(0.5)AVTYDSPYPLYAMAFSSLR* | *S(0.855)VVET(0.145)QLIAHDKEVHDIAWGEAR* | *S(0.093)ET(0.907)AVTYDSPYPLYAMAFSSLR* | *S(0.375)VVET(0.625)QLIAHDKEVHDIAWGEAR* |
| **Identification type KIN10K48M-TTG1** | | | | | | | *By MS/MS* |
| **Identification type KIN10-TTG1** | *By MS/MS* | *By MS/MS* | *By MS/MS* | *By MS/MS* | *By MS/MS* | *By MS/MS* | *By matching* |
| **Intensity** | *418430000* | *155640000* | *249750000* | *3E+07* | *9757500* | *4,8E+07* | *3,9E+07* |
| **Intensity___1** | *418430000* | *155640000* | *249750000* | *3E+07* | *9757500* | *4,8E+07* | *3,9E+07* |
| **Intensity___2** | *0* | *0* | *0* | *0* | *0* | *0* | *0* |
| **Intensity___3** | *0* | *0* | *0* | *0* | *0* | *0* | *0* |
| **Ratio mod/base** | *0.017591* | *0.0065431* | *0.0105* | *0.0020532* | *0.00033808* | *0.0032765* | *0.0013388* |
| **Intensity KIN10K48M-TTG1** | *0* | *0* | *0* | *0* | *0* | *0* | *2,6E+07* |
| **Intensity KIN10-TTG1** | *79589000* | *155640000* | *166640000* | *3E+07* | *9757500* | *3E+07* | *1,2E+07* |
| **Ratio mod/base KIN10K48M-TTG1** | *0* | *0* | *0* | *0* | *0* | *0* | *0.0016041* |
| **Ratio mod/base KIN10-TTG1** | *0.0093335* | *0.018252* | *0.019542* | *0.0055354* | *0.0007787* | *0.0055354* | *0.00099366* |
| **Intensity KIN10K48M-TTG1___1** | *0* | *0* | *0* | *0* | *0* | *0* | *2,6E+07* |
| **Intensity KIN10K48M-TTG1___2** | *0* | *0* | *0* | *0* | *0* | *0* | *0* |

| | | | | | | | |
|---|---|---|---|---|---|---|---|
| ***Intensity KIN10K48M-TTG1___3*** | *0* | *0* | *0* | *0* | *0* | *0* | *0* |
| ***Intensity KIN10-TTG1___1*** | *79589000* | *155640000* | *166640000* | *3E+07* | *9757500* | *3E+07* | *1,2E+07* |
| ***Intensity KIN10-TTG1___2*** | *0* | *0* | *0* | *0* | *0* | *0* | *0* |
| ***Intensity KIN10-TTG1___3*** | *0* | *0* | *0* | *0* | *0* | *0* | *0* |
